# Dopamine and GPCR-mediated modulation of DN1 clock neurons gates the circadian timing of sleep

**DOI:** 10.1101/2021.12.16.472997

**Authors:** M. Schlichting, S. Richhariya, N. Herndon, D. Ma, J. Xin, W. Lenh, K. Abruzzi, M. Rosbash

## Abstract

The metronome-like circadian regulation of sleep timing must still adapt to an uncertain environment. Recent studies in *Drosophila* indicate that neuromodulation not only plays a key role in clock neuron synchronization but also affects interactions between the clock network and brain sleep centers. We show here that the targets of neuromodulators, G-Protein Coupled Receptors (GPCRs), are highly enriched in the fly brain circadian clock network. Single cell sequencing indicates that they are not only differentially expressed but also define clock neuron identity. We generated a comprehensive guide library to mutagenize individual GPCRs in specific neurons and verified the strategy with a targeted sequencing approach. Combined with a behavioral screen, the mutagenesis strategy revealed a novel role of dopamine in sleep regulation by identifying two dopamine receptors and a clock neuron subpopulation that gate the timing of sleep.

## Introduction

Animal function requires communication between different brain centers. An important part of this communication is neurotransmission, which takes place at synapses and leads to the excitation and/or inhibition of downstream neurons. Recent studies stress the importance of a second means of communication, namely neuromodulation, in the central nervous system. In contrast to classical neurotransmission, neuromodulators do not directly lead to the opening of ion channels but alter second messengers, which then affect electrophysiological responsiveness ^1,2^. They can also influence transcriptional activity and metabolism. Although neuromodulation is not a new concept, several key issues are not yet well-understood. Among them is the network organization of their target receptors.

All neuromodulators interact with G-Protein Coupled Receptors (GPCRs) ^3^. This class of receptors has seven transmembrane domains, are located within the plasma membrane and react to a variety of stimuli, including neuropeptides, biogenic amines and even light. In an inactive state, many receptors are coupled to a heterotrimeric G-protein consisting of alpha, beta and gamma subunits. Depending on the identity of the alpha subunit, these proteins dissociate upon receptor activation and lead to increases in Ca^2+^, cAMP or to the activation of transcription via the rho pathway ^3,4^. GPCRs are therefore useful for the study of brain circuits, as they directly influence the molecular properties of individual neurons.

The circadian clock neuron network (CCNN) of *Drosophila melanogaster* is an ideal model to study the contribution of GPCR signaling to an adult brain circuit. The fly brain CCNN only consists of 150 neurons, and diverse modes of interactions and distinct functions have been assigned to specific cells and several studies emphasize the abundance of circadian neuropeptides and their importance to the functions of this network ^5–10^. Pigment dispersing factor (PDF) is a key clock neuropeptide and is expressed in two sets of lateral neurons, the 4 large-ventrolateral neurons (lLNv) and the 4 small ventrolateral-neurons (sLNv) ^11^; they are important for arousal and morning (M) activity, respectively ^12–15^. PDF is believed to synchronize its downstream targets, and a loss of the peptide leads to arrhythmicity in constant darkness (DD) ^5,16^. Three of the six dorso-lateral neurons (LNd) and the 5th sLNv serve a different function and are important for evening (E) activity ^12,13^. A pair of dorsal neurons (DN2s) are essential for temperature preference rhythms ^17^. No function has been assigned to the DN3s, perhaps due to the lack of a specific driver. The DN3s are the most numerous clock neuron group (35-40 neurons/hemisphere).

The second most numerous clock neuron group is the DN1 cluster, which consists of approximately 15 neurons per hemisphere ^17^. Accelerating or decelerating the clock in these cells has no effect on rhythmic behavior under DD (constant darkness) conditions, and the DN1s are even dispensable for rhythms in DD ^18–21^. However, changing the speed of these cells shifts the timing of the evening (E) activity peak in a LD (light:dark) cycle, indicating a DN1p role in circadian timing.

Moreover, several groups have shown that specific DN1p neurons affect morning (M) as well as E activity and influence fly sleep, both the amount of sleep and when sleep occurs ^22–24^. Consistent with a role in sleep promotion and/or maintenance, imaging and tracing experiments identified physiological pathways connecting specific DN1ps to fly brain sleep centers ^22^.

Despite general agreement on the variety of functions carried out by DN1ps, there are discrepancies in assigning specific behavioral roles to specific DN1p subgroups. For example, glutamatergic DN1ps have been identified as controlling the morning component of behavior, whereas another study suggests that these same neurons control E activity ^23,24^. A likely explanation is that the DN1ps are even more diverse than previously thought, i.e., there may be multiple glutamatergic DN1p subgroups. Indeed, a recent single cell seq study found that the DN1ps are composed of at least six different clusters, consistent with the notion that different DN1p functions might derive from different neuron subpopulations ^25^.

To address the contribution of DN1ps and neuromodulation to sleep behavior, we combined the power of single-cell RNAseq (scRNAseq) data with cell-specific CRISPR/Cas9 mediated gene mutagenesis. The scRNAseq results not only show that GPCRs are highly enriched in the CCNN but also that they are key components for defining clock neuron identity. To mutate specific receptors in a cell-specific manner, we generated a CRISPR/Cas9 based guide library for all GPCRs following the pioneering work of Port and Bullock^26^. We and others had previously demonstrated that this strategy efficiently removes PER or TIM expression from the clock network in a cell-specific manner ^27,28^. Here we also added an adapted targeted genomic sequencing approach to show that the library effectively mutates GPCRs in a cell-specific manner, suggesting that the library and strategy constitutes a key asset tool for investigating neuromodulation. Indeed, we used the library in a behavioral screen that identified several GPCRs that promote sleep from within the CCNN. In addition to already known sleep-promoting GPCRs, two dopamine receptors (Dop1R1 and Dop1R2) promote sleep via the DN1ps. Combining state-of-the-art tracing techniques with single cell data moreover shows that different subsets of DN1ps contribute differentially to sleep. These data taken together indicate that dopamine regulates sleep timing via a novel pathway through the CCNN and suggests that its actions are highly context-specific. These findings and methods will facilitate investigating the complex contribution of dopamine as well as other neuromodulators and neurotransmitters in various other physiological processes.

## Results

### Circadian neurons show enriched expression of signaling molecules

Recent work implicates clock neuron interactions in normal circadian behavior. For example, manipulating clock neuron subpopulations can affect behavioral timing, and residual clock protein expression in only a few circadian neurons is sufficient to retain some circadian function ^9,21,27–29^. To identify molecules within the clock network that contribute to the implied network synchrony, we isolated and sequenced FACS-sorted clock neurons (*clk856>EGFP*) under LD conditions at two times, ZT2 and ZT14, and compared the transcriptomes to those from all neurons (*nSyb>EGFP*).

We first compared clock gene expression between the time points in the clock neurons. Consistent with expectation ^30^, *clk* mRNA levels were approximately 6x higher at ZT2 than at ZT14, whereas *tim* mRNA levels were 22x higher at ZT14 compared to ZT2 (Fig. 1A). Housekeeping genes such as *Act5C* and *Rpl32* were not different between time points (Fig. 1A). Not surprisingly, clock gene expression was dramatically reduced in the *nSyb* dataset (*clk*: 0.07%; *tim*: 3.7%, Fig. S1), demonstrating highly efficient enrichment of clock neurons by *clk856>EGFP* purification.

**Figure 1.**
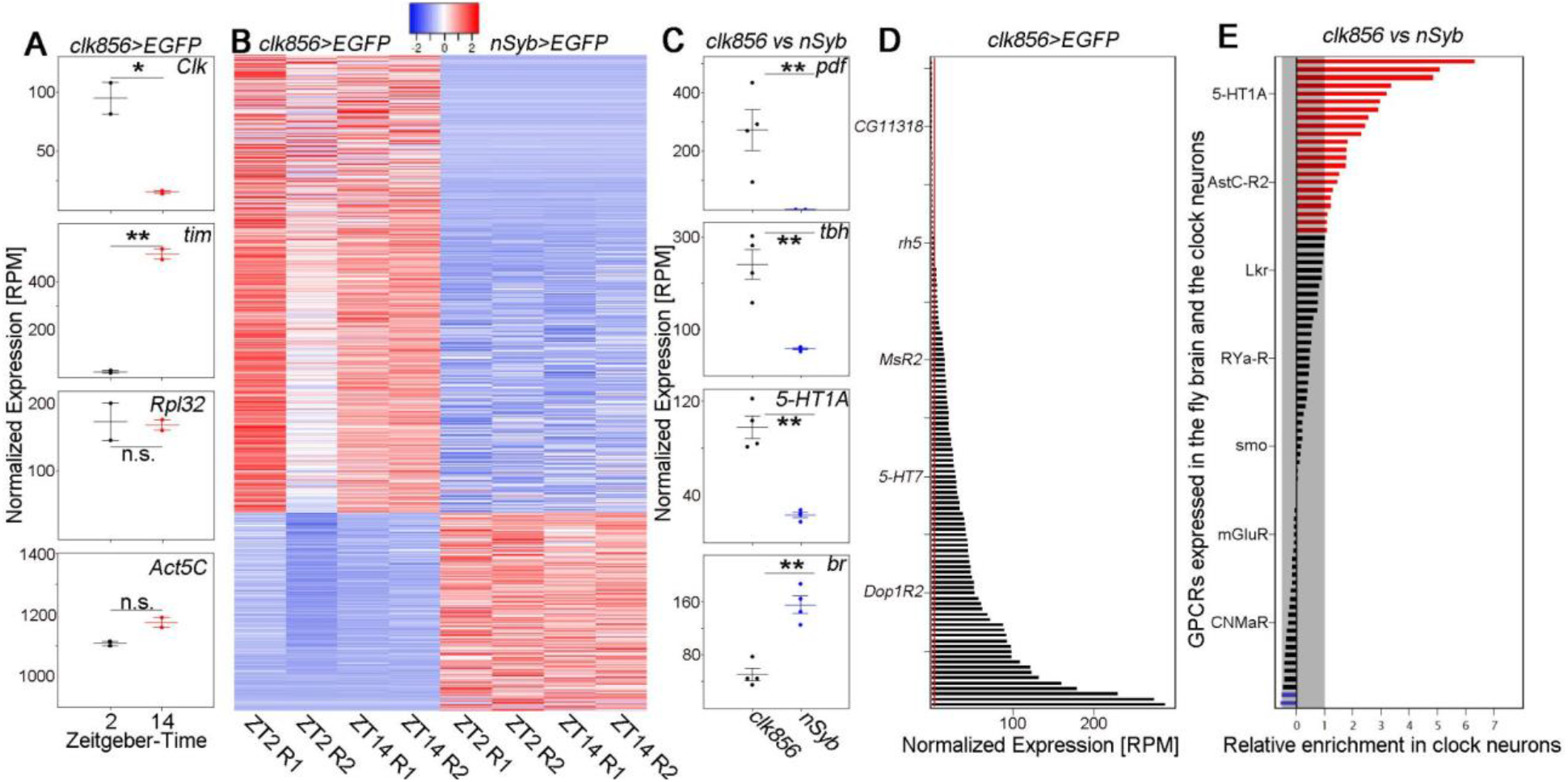
Clock neurons show enriched GPCR expression. **A** Plots show normalized expression levels of indicated genes from sorted clock neurons (*clk856>EGFP*) at ZT2 and ZT14. As expected, *Clk* expression is significantly higher at ZT2 than at ZT14, whereas *tim* expression is significantly higher at ZT14 compared to ZT2. Housekeeping genes such as *Rpl32* and *Act5C* were not significantly different between those timepoints. **B** Heatmap of differentially expressed genes between clock neurons (*clk856>EGFP*) and randomly chosen neurons (*nSyb>EGFP*) at ZT2 and ZT14 with an at least 2-fold difference in expression at FDR<0.05. Relatively high expression is displayed in red and relatively low expression is displayed in blue (Z-scores indicated on top of the graph). **C** Examples of up- or downregulated genes representing enriched GO terms. *Pdf* is significantly upregulated in the clock network (GO: circadian control of sleep wake cycle) as are *tbh* (GO: octopamine and tyramine signaling pathway) and *5-HT1A* (GO: G-protein coupled receptors). *Br* on the other hand is significantly downregulated in the clock network (GO: Antennal development). **D** Normalized expression (RPM) of GPCR genes in the clock network. GPCR gene-expression varies dramatically between receptors from being not expressed (< 3RPM, red line) to being highly expressed. **E** Relative expression of GPCRs in clock neurons (*clk856>EGFP*) relative to randomly chosen neurons (*nSyb>EGFP*). 22 GPRs are at least 2-fold higher expressed in the clock cells (red bars) compared to only 2 GPCRs being at least 2-fold downregulated in the clock cells (blue bars).

To address clock neuron gene enrichment more generally, we used edgeR to perform differential gene expression analysis. It identified a total of 1719 genes as differentially expressed with false discovery rate <0.05 between *nSyb* and clock neurons. Of these, 704 genes were significantly upregulated and 305 genes were significantly downregulated in the clock network with at least a 2-fold change in amplitude (Fig. 1B). GO-term analysis of clock-enriched genes resulted in enrichment in expected categories like the circadian regulation of temperature homeostasis (27-fold), positive regulation of circadian sleep/wake cycle (16-fold) and circadian regulation of gene expression (11-fold). Interestingly, genes associated with signaling pathways were also found to be comparably enriched among the clock-enriched genes and include the octopamine and tyramine signaling pathways (16-fold), the serotonin receptor signaling pathway (16-fold) and the G protein-coupled receptor signaling pathway (12-fold). Other unrelated pathways are enriched in the *nSyb* neurons relative to the clock neurons (Fig. 1C). These data strongly implicate neuronal communication in clock network function.

### GPCRs are differentially expressed within the clock neuron network

The enrichment of GPCRs inspired a focus on quantitative features of their expression. Surprisingly, more than 2/3 of the 124 GPCR mRNAs encoded by the *Drosophila* genome are present in the clock neurons (Fig. 1D) with an almost 100x difference between the least and most expressed of these 86 mRNAs. Similar trends were also found in our *nSyb* data (Fig. S2). A direct comparison with the *nSyb* dataset indicates that 22 GPCRs are at least 2-fold upregulated in the clock network, whereas only 2 receptors are downregulated, further supporting the importance of neuromodulation to the clock network (Fig. 1E).

The expression data suggest that GPCRs are expressed at different levels in all clock neurons or that they are predominantly expressed in neuron subpopulations. To distinguish between these possibilities, we analyzed previous single cell RNAseq data for GPCR expression and focused on the 17 high confidence clock neuron clusters (Fig 2A); they are missing most of the enigmatic DN3 clock neurons but include most if not all well-characterized clock neurons including all lateral and most dorsal clock neurons.

**Figure 2.**
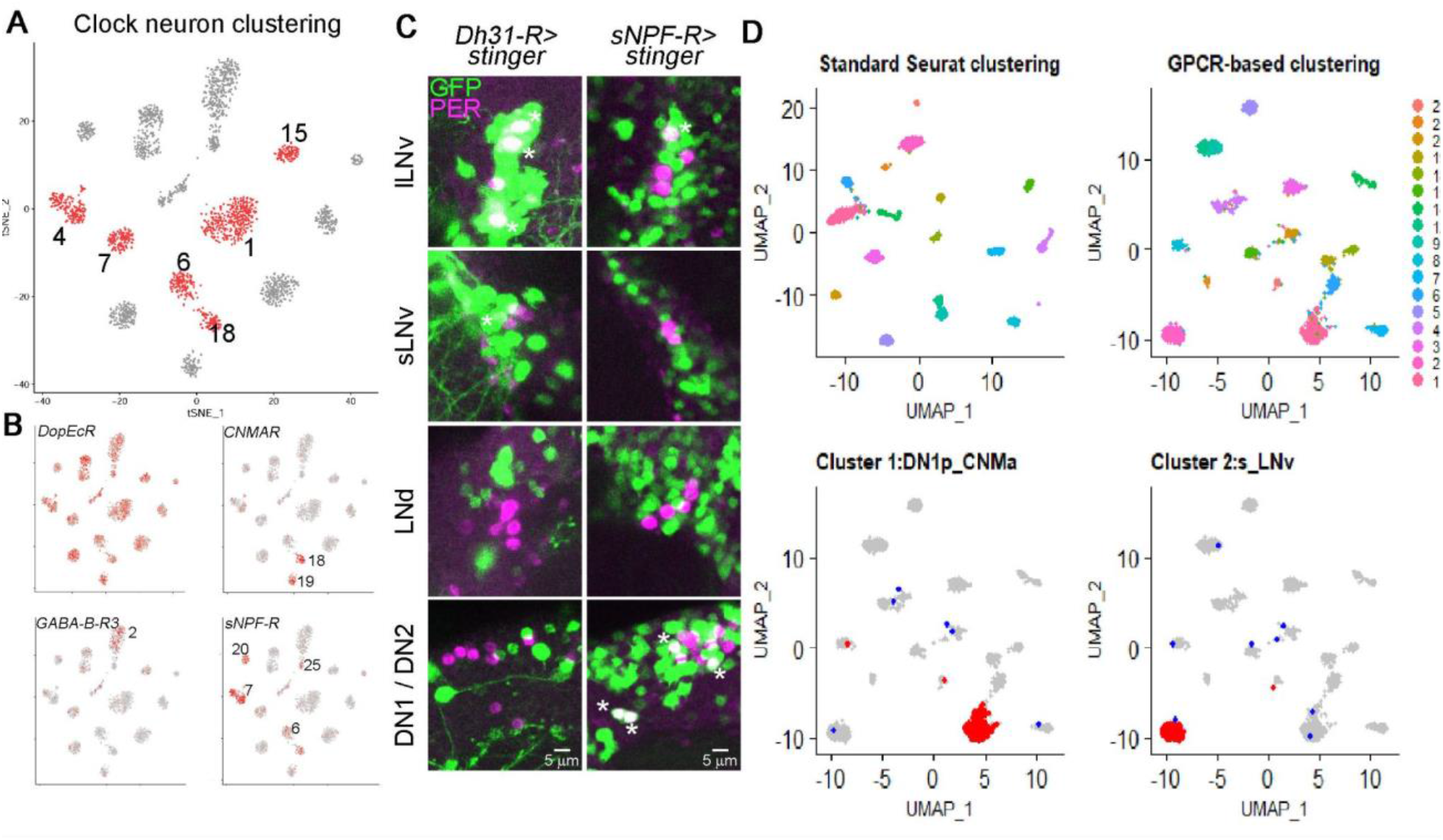
GPCRs are differentially expressed and can define clock neuron identity. **A** Single cell RNA sequencing of clock neurons (*clk856>EGFP*) identifies 17 bona-fide clock neuron clusters. DN1 neurons can be separated into six different clusters (1, 4, 6, 7, 15, 18, labeled in red). For details on clustering see ^25^ . **B** Expression of different GPCRs within these 17 clusters. *DopEcR* is highly expressed in all clock neuron clusters, whereas *CNMaR, GABA-B-R3* and *sNPF-R* are more differentially expressed. For details see text. **C** Immunohistochemistry of whole mount brains stained against GFP (green) and PER (magenta). Nuclear GFP (UAS-*stinger*) was expressed under the control of endogenous GAL4 lines. *Dh31-R-GAL4* drives expression in 3 out of 4 lLNvs, 1 sLNv and does not express in DN1s, DN2 or LNds. *sNPFR-GAL4* is expressed in 1 lLNv, 2 DN1s and the DN2 neurons. **D** Seurat clustering of individual neurons included in 17 high-confidence clusters. Unsupervised clustering (top left) recapitulates previously published results. Similarly, clustering only based on GPCR expression generated 17 different clusters (top right). Bottom row: Analysis of retained (red) or newly assigned (blue) clock neuron identities when plotted only based on GPCRs. Most neurons in clusters 1 (bottom left) and cluster 2 (bottom right) are assigned the same clusters (red dots) whereas only a few cells were assigned to different clusters (blue dots) based on GPCR-only clustering of clock cells.

The dopamine receptor *DopEcR* is highly expressed in the *nSyb* as well as the *clk856* dataset with high transcript levels in all clock neuron clusters (Fig. 2B). We then analyzed the expression patterns of more poorly expressed GPCRs and found impressive differential expression within the clock neuron population. For example, *CNMaR* appears to be almost exclusively expressed in one DN1p and the DN2 cluster, whereas *Gaba-B-R3* is highly enriched in the s-LNv cluster. sNPFR is expressed more broadly but still mostly detected in the clusters defining the DN1ps, the lLNvs and the DN3s (Fig. 2B). A differential GPCR expression pattern is also apparent when comparing all GPCR expression using single-cell RNAseq despite dramatic variation in expression levels (Fig. S3).

To confirm this cell-specificity in an independent way, we used GAL4 lines in which the GAL4 sequence was integrated into endogenous GPCR-expressing loci and used to express nuclear GFP (*UAS-stinger*) ^31^. Notably, the overall expression levels of individual GPCRs correlated nicely with the *nSyb* sequencing data (Fig. S2). Some GPCRs such as the aforementioned *DopEcR* appear to be expressed almost pan-neuronally within the brain, whereas others are not expressed in the brain or only in 1-2 cells per hemisphere (Fig. S2).

To detect the expression of specific GPCRs within the clock neuron network, we co-stained with anti-PER and determined the overlap of PER and GFP. We also focused on GPCRs with striking expression patterns from our single cell data. *DH31-R* appears highly enriched in the lLNvs and slightly enriched in the sLNvs and the DN1ps. The endogenous GAL4 line is strikingly consistent with this pattern: 3 out of 4 lLNvs were GFP-positive, whereas only 1 out of 4 sLNv neurons was labeled, explaining the differences in expression levels. As predicted, no LNds were labeled, but we could also not detect GFP expression in the DN1 neurons in most brains. Similar cell-type specificity was also observed with *sNPFR*, which mainly labeled DN1 neurons and 1 out of 4 lLNvs (Fig. 2C). The data taken together suggest that GPCRs are strongly differentially expressed within the clock network. Notably, differences appear even between individual cells within a supposed uniform cluster like the lLNvs, as often not all cells express a GPCR of interest (Fig. S4).

These striking differences in GPCR expression suggested that these receptors might be a defining feature of clock cell identity within the clock network. To address this question, we first used the single cell data and a standard seurat clustering method, which was able to identify all 17 previously identified clusters (Fig. 2D). We then only used differences in GPCR expression and could still identify 17 distinct clusters (Fig. 2D). To our surprise, many of these newly generated clusters could be mapped onto the previously published dataset ^25^, indicating that the GPCR-generated clusters are of biological and anatomical significance. For example, the newly identified cluster 1 includes approximately 90% of the previously identified cluster, which represents the sLNvs. This is also the case for other clock clusters like the DN1s (Fig. 2D). The data show that GPCR expression is sufficient to specify clock neuron identity.

### A guide library allows for cell-specific manipulations of all GPCRs

The likely importance of GPCRs for clock neuron identity and the requirement of neuropeptides for clock synchrony inspired the development of a general strategy to eliminate any GPCR in a neuron-specific manner. Relevant to this goal, we and others recently showed that CRISPR/Cas9 based mutagenesis is superior to RNAi-mediated gene expression knockdown in the fly brain ^27,28^. As the former also does not require strongly expressing driver lines, it is much more amenable to highly neuron-specific split-GAL4 lines.

The *Drosophila* genome encodes 124 GPCRs, which are known to react to a variety of stimuli, including biogenic amines, neurotransmitters, neuropeptides and even light (Fig 3A) ^4^. To manipulate any of these receptors, we generated UAS-guide lines, which express three guides targeting the coding sequence of a GPCR. Three guides have previously been shown to efficiently mutate eye tissue and provide high mutagenesis efficiency by compensating for potential non-cutting guides ^26^; this strategy also worked well to remove PER and TIM from the clock system ^27,28^. As there are no reliable antibodies for many of the receptors, we designed a targeted genomic sequencing approach to verify the functionality of the library. We used *clk856*-GAL4 to drive the expression of GFP, Cas9 and the guide of interest in most of the clock network. The clock neurons are labeled with GFP and simultaneously mutated by the CRISPR/Cas9 system. We then dissected and dissociated the fly brains, FACS-sorted the neurons and extracted their DNA.

**Figure 3.**
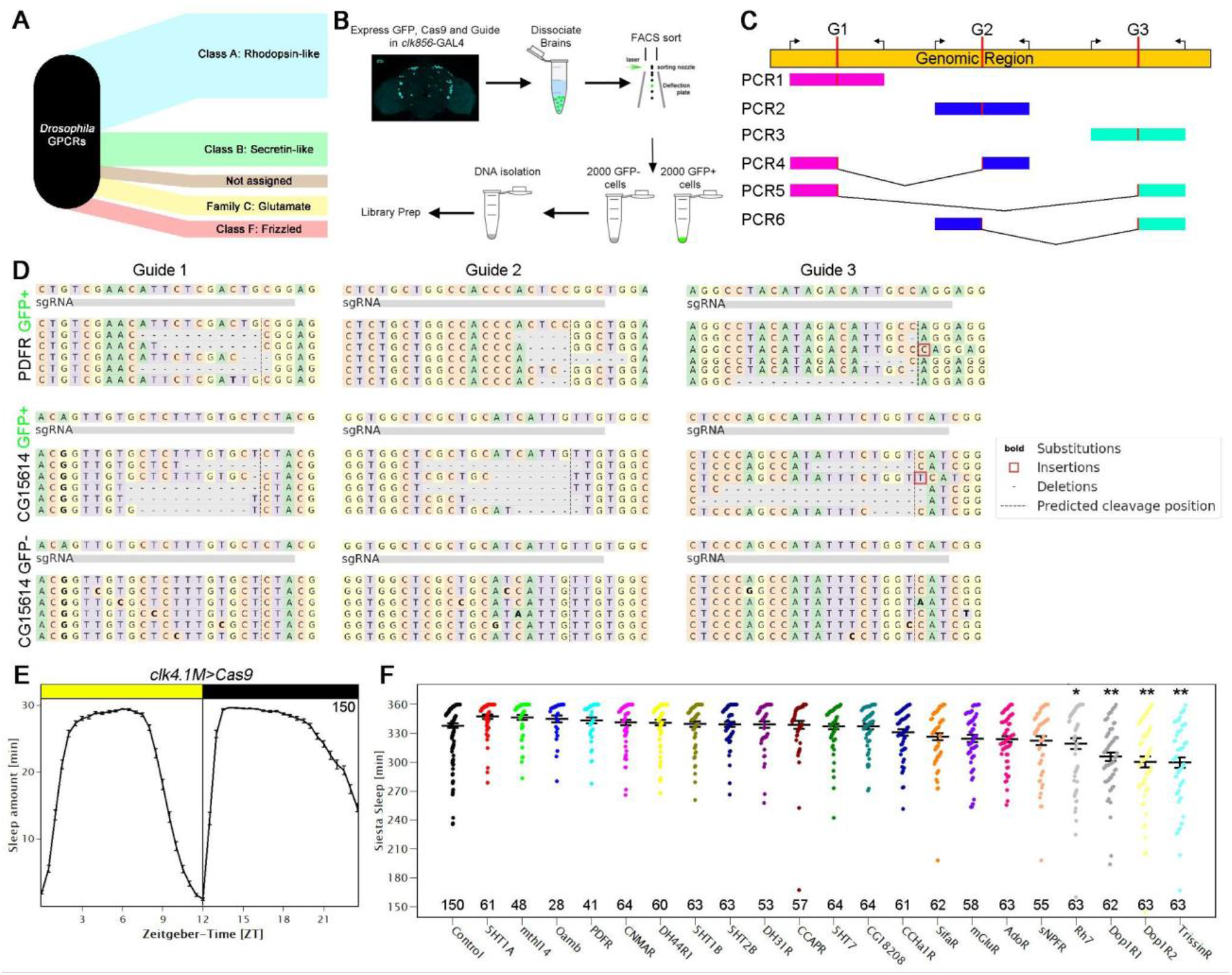
A guide-RNA library for all GPCRs of the fly genome allows for cell specific mutagenesis. **A** *Drosophila* GPCRs can be categorized into 5 distinct groups (reviewed in ^4^) **B** Workflow of the targeted genomic sequencing assay. Clock neurons were labeled with EGFP (*clk856>EGFP*) while expressing Cas9 and the guide-RNA of interest which allows for cell-specific manipulations. GFP positive cells were analyzed for possible mutagenesis. GFP negative cells were used as controls for cell specificity. **C** Schematic of genomic sequencing approach. Each genomic region was targeted by three independent guide-RNAs (G1-G3) which lead to double-strand breaks (DSB) at the desired locations (indicated in red). We simultaneously used three sets of primers for each reaction amplifying approximately 250 bp. Depending on the timing of guide-RNA mediated DSBs, we expect either short deletions (PCR1-3) or big deletions (PCR4-6) in the event of two guides cutting at the same time. **D** Genomic sequencing results of targeted genomic sequencing. Top panel: Sequences of GFP-positive cells of *clk856>EGFP, Cas9, PDFR-g* flies. All three guides are able to cause small deletions at the expected site of DSB. Middle panel: Sequences of GFP-positive cells of *clk856>EGFP, Cas9, CG15614-g* flies. All three guides are able to cause small deletions at the expected site of DSB. Lower panel: Sequences of GFP-negative cells of *clk856>EGFP, Cas9, CG15614-g* flies. No deletions were detected in GFP-negative cells. **E** Sleep behavior of control flies ± SEM expressing *Cas9* in DN1 clock neurons (*clk4*.*1M>Cas9*). The flies showed the expected sleep pattern with low sleep in the morning and evening and high levels of sleep at night and during the siesta indicating that expressing Cas9 in the dorsal neurons does not affect normal sleep behavior. **F** Sleep amount of flies with mutated GPCRs during the siesta (ZT3-9) in LD. *Clk4*.*1M>Cas9* flies were used as controls, for experimental flies *Clk4*.*1M>Cas9* flies were crossed to UAS-guide-RNA lines as indicated on the x-axis. Four guide-RNAs significantly reduced sleep during the siesta. Numbers indicate the number of flies used for the behavioral screen. * p<0.05 ** p<0.01.

To address the question of cell-specificity, we sorted 2000 GFP-positive cells, which should have been mutated, and 2000 GFP-negative cells, which should remain wild-type (Fig. 3B). We designed three sets of primers flanking each of the guide binding sites to allow for gene-specific amplification. As three guides are being used at the same time, there can either be small deletions in the area of guide-binding (Fig. 3C, PCR 1-3) or larger deletions if multiple guides cut at the same time, resulting in different DNA fragment combinations (Fig. 3C, PCR 4-6). We also analyzed five different guide strains on three different chromosomes (*PDFR* and *Tre1* on the X-chromosome, *mAchR-A* and *CG15614* on the 2nd chromosome and *CrzR* on the third chromosome) to avoid possible biases from chromosome location. *PDFR* served as a positive control: guide-mediated mutagenesis of the clock network with its guides completely reproduced *PDFR* full body mutant (*han*^*5304*^) phenotypes (data not shown) ^32,33^.

We first analyzed the libraries of the GFP-positive cells and found that all three *PDFR* guides were able to generate deletions of variable sizes at the expected cut sites (Fig. 3D). Moreover, we were able to generate large deletions as we obtained several thousand reads for a genomic fragment representing a deletion of several thousand base pairs (Fig. 3C, PCR5). GFP-negative cells in contrast showed no deletions in the investigated area, suggesting that there is no background mutagenesis due to leaky expression in non-target cells. We then assayed *CG15614*. Like for *PDFR*, all three guides generated deletions, whereas GFP-negative cells were unaffected (Fig. 3C). *Tre1* and *CrzR* had similar results (data not shown), whereas only two out of the three guides for mAchR-A created deletions (Fig. S5).

### DN1p modulation alters the sleep structure of male flies

To exploit this functional library, we modified GPCR expression in subsets of clock neurons and focused on the DN1ps. They were previously shown to affect sleep and connect to deep-brain sleep centers ^22–24^. We used *clk4*.*1M*-GAL4, which expresses in 8-12 of the 15 DN1ps per hemisphere. We first reproduced previous experiments: activating these neurons significantly altered siesta sleep of male flies (Fig. S6).

To identify candidate GPCRs, we turned to our single cell data and identified 21 GPCRs, which are enriched in DN1ps compared to the other clock cells. We then performed a behavioral screen in which we compared the behavior of flies with mutated GPCRs in DN1ps compared to control flies only expressing *Cas9* in the DN1s (*clk4*.*1M>Cas9*). As expected, *Cas9* expression in the DN1ps did not affect fly behavior: they showed the canonical sleep pattern with consolidated sleep at night and during the siesta with almost no sleep in the morning and the evening; this reflects the standard bimodal activity pattern (Fig. 3E). We then focused on the siesta and quantified sleep levels between ZT3 and ZT9. Male flies sleep extensively during this time, leading to a median of approximately 5.5 hours of sleep in control flies. Of the 21 mutated strains, four significantly reduced their siesta sleep compared to the control group by one-way Anova followed by a post-hoc Tukey test (Fig. 3F). One GPCR is *rh7*, which reproduces known whole-body mutant phenotypes ^34^. Another is *TrissinR*, which reduced sleep by half an hour. Trissin was previously found to be expressed in 2 LNds, suggesting that intra-clock neuron communication, LNd to DN1p, is relevant to sleep regulation ^25^. Eliminating two different dopamine receptors, Dop1R1 and Dop1R2 also reduced siesta sleep.

### Dopaminergic input to DN1 neurons reduces siesta sleep

The fact that mutating Dop1R1 and Dop1R2 in the DN1ps reduced sleep was very surprising, as dopamine is traditionally activity-promoting ^35–37^. We therefore addressed siesta sleep more specifically by dividing the 24 hour day into 4 equal sections of 6 hours each: ZT21-3 for morning sleep, ZT3-9 for siesta sleep, ZT9-15 for evening sleep and ZT15-21 for nighttime sleep (Fig. 4A). We first quantified sleep with *Dop1R2* mutated in DN1ps. Although sleep levels in the mutated flies were dramatically different in the 4 different intervals as expected (Fig. 4B), sleep was only significantly reduced during the siesta (p<0.01) compared to the Cas9 (*clk4*.*1M>Cas9*) and Guide (*clk4*.*1M>Dop1R2-g)* control flies. Very similar effects were found when mutating *Dop1R1* in the DN1ps (Fig. S7).

**Figure 4:**
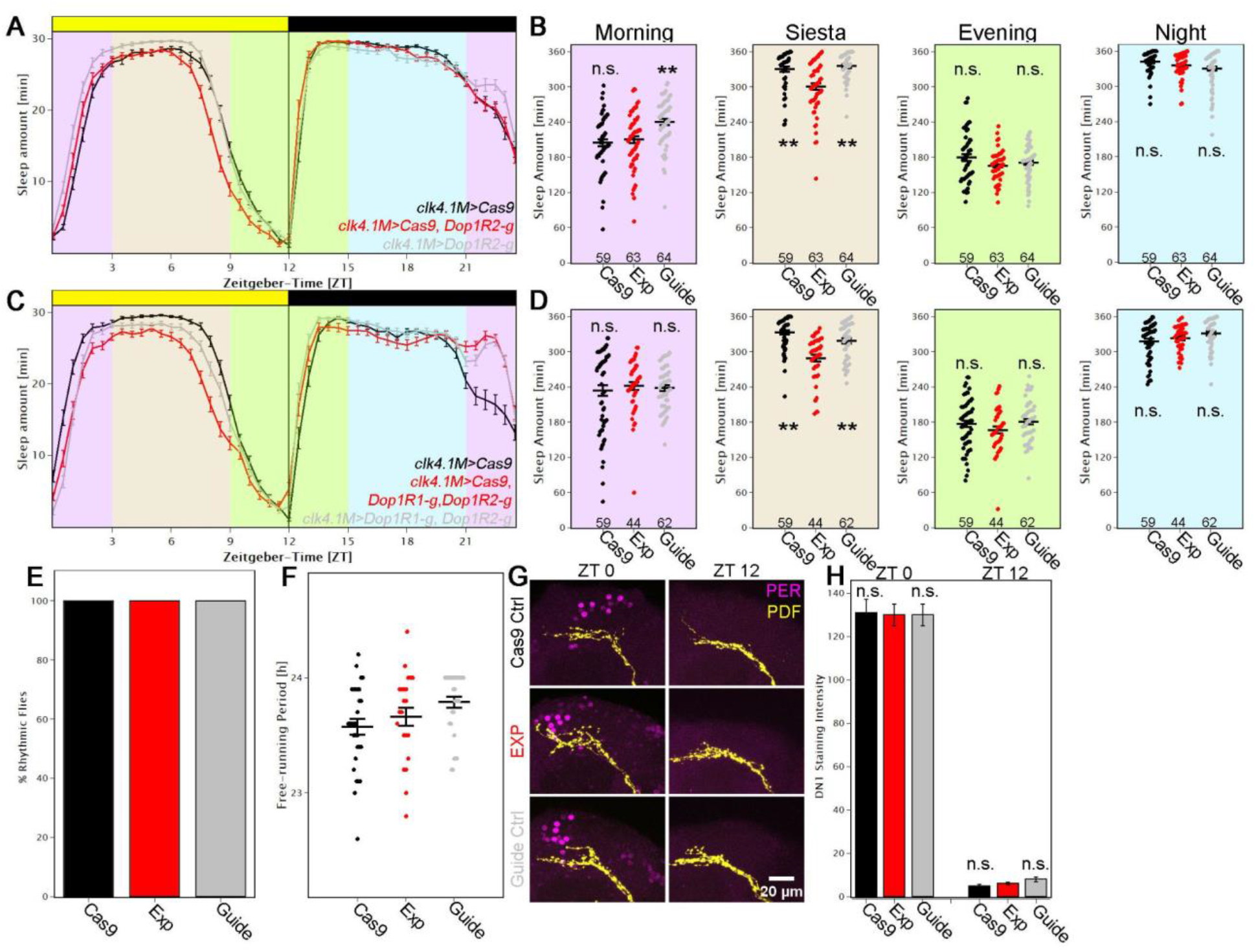
Modulation of DN1ps by dopamine enhances sleep. **A** Average sleep profile of male flies in which *Dop1R2* was mutated in the DN1ps (red) and controls (grey and black). Error bars represent SEM. Background colors indicate 4×6h periods which were quantified in B. **B** Quantification of sleep separated into 4 different time zones: Morning (ZT21-ZT3), Siesta (ZT3-ZT9), Evening (ZT9-ZT15) and Night (ZT15-ZT21). Removing *Dop1R2* in the DN1ps significantly reduced sleep during the siesta (p<0.01), whereas other times of day were not affected. **C** Average sleep profile of male flies in which *Dop1R2* and *Dop1R1* were mutated in the DN1ps (red) and controls (grey and black). Error bars represent SEM. Background colors indicate 4×6h periods which were quantified in D. **D** Quantification of sleep separated into 4 different time zones). Removing *Dop1R2* and *Dop1R1* in the DN1ps significantly reduced sleep during the siesta (p<0.01), whereas other times of day were not affected. **E-F** Flies with mutated *Dop1R1* and *Dop1R2* in the DN1ps (EXP) have rhythmicity same as controls (**E**) and no effect in free-running period (**F**) when recorded in DD compared to both controls (*clk4*.*1M>Cas9* (black) and c*lk4*.*1M>DopiR1-g, Dop1R2-g* (grey)). **G** Flies with mutated *Dop1R1* and *Dop1R2* in the DN1ps (EXP) and both controls (*clk4*.*1M>Cas9* (black) and c*lk4*.*1M>DopiR1-g, Dop1R2-g* (grey)) stained with anti-PDF and anti-PER at ZT0 and ZT12. All brains show the expected high levels of PER at ZT0 and low levels at ZT12 as quantified in **H**.

To provide further reassurance that dopamine is sleep-promoting in the DN1ps, we expressed *Cas9* and guides against *Dop1R2* in the dorsal fan-shaped body and could reproduce the increased siesta sleep phenotype observed by Pimentel et al ^38^ using the same dFSB driver *23E10* and RNAi knockdown of *Dop1R2* (Fig. S8). This result indicates that the target receptors and cells dictate the effect of dopamine, which promotes sleep as well as wake.

Will mutating both *Dop1R1* and *Dop1R2* at the same time have an even stronger effect on sleep? Although siesta sleep is significantly reduced compared to both control strains with no effect at any other time of day, we did not observe any additive effect: the double-mutated strain had a similar effect compared to the single mutant strains. (Fig. 4C-D).

These sleep patterns indicate that the siesta is terminated earlier in flies with mutated dopamine receptors compared to both controls, suggesting that dopaminergic input contributes to timing the end of the siesta. Notable in this context is the traditional *per*^*S*^ mutant strain: it has a very short free-running period with a similar LD phenotype, e.g., the E peak occurs during the daytime ^38,39^. Yet there was no effect of removing both dopamine receptors in the DN1ps on period length and rhythmicity: experimental groups as well as controls were identical to wild-type (Fig 4E, F), suggesting that changes in clock speed are not responsible for the sleep phenotypes. Mutating the dopamine receptors also had no visible effects on PER expression: PER antibody staining indicated high and normal PER levels at ZT0 and low levels at ZT12 (Fig. 4G-H). The results taken together indicate that there are no major effects of dopamine receptor removal on rhythmicity or clock protein expression, which supports the notion that dopamine gates the timing of sleep by modulating dorsal neuron function under LD conditions.

### Sub-clustering of DN1 neurons allows to differentiate neurons controlling E activity

Can the mutant dopamine receptor phenotype help characterize the clock neurons that influence E activity? To this end, we used the trans-tango technique to identify target neurons downstream of the dopaminergic system. Because large portions of the brain are labeled, we co-labeled the brains with anti-PER and investigated co-localization of GFP and PER immunoreactivity. The dopaminergic system appears to contact a variety of clock cells including PDF cells, 3 out of 6 LNds, DN2s and on average 4 DN1 neurons, confirming that the DN1s are indeed direct downstream partners of the TH cells (Fig. S9).

Examining the six DN1 clusters in more detail indicated that *Dop1R1* and *Dop1R2* are primarily expressed in clusters 6, 7, 15 and 18 (Fig. 5A). Intriguingly, this number of clusters, 4, is identical to the number of TH neuron target DN1ps. Importantly, the size of the individual clusters is also rather small, suggesting that several of these clusters contain a single cell ^25^. We also considered the two remaining DN1p clusters, clusters 1 and 4. The biggest cluster (#1) is the only one that expresses *AstC*. We showed previously that *AstC* is expressed in four DN1 neurons, which fits well with the size of this cluster ^8,25^. In addition, knockdown of *AstC* affected the timing of the E peak in summer or winter days, suggesting that these neurons also contribute to E activity at least in different seasons. In addition, this is the only cluster expressing *TrissinR* which also produced a siesta phenotype in our behavioral screen (Fig. S7). These data taken together indicate that several if not most DN1 clusters contribute to E activity. The exception is cluster 4, which by elimination probably accounts for prior results indicating that the DN1s also affect M activity ^40^.

**Figure 5.**
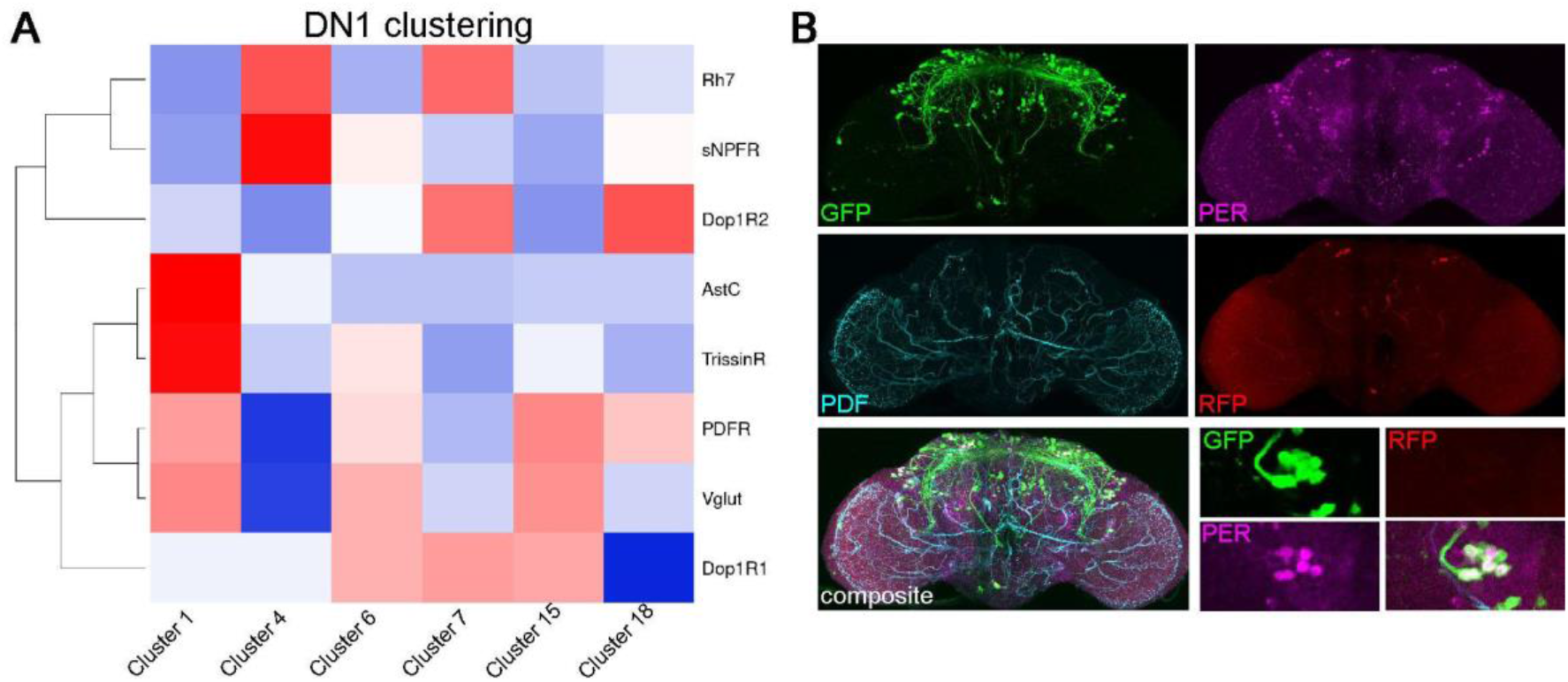
Sub-clustering of DN1ps indicates their function in controlling E activity. **A** Clustering of DN1ps using key molecules expressed in these cells. **B** Trans-tango experiment of the *Vglut-DBD, per-AD* split-GAL4 driver line stained with anti-GFP (trans-tango positive neurons), anti-PDF, anti-PER and anti-RFP (GAL4 expressing neurons). Several neurons in the dorsal and lateral part of the brain are downstream of the *Vglut*-positive DN1 neurons. Notably, all LNds are included indicating an anatomical connection of siesta behavior with cells controlling E activity. For details see text.

How do these DN1s influence the siesta and E activity? A strategy to answer this question can exploit *Vglut* expression. It is expressed in most DN1 clusters affecting siesta or E activity but not in cluster 4 (Fig. 5A). Combining a *per*-AD with *Vglut*-DBD labeled on average 7-8 DN1s per hemisphere, consistent with the expected number of neurons in these clusters (Fig. S10) Trans-tango experiments with this split-GAL4 revealed that these neurons primarily target neurons in the dorsal part of the brain (Fig. 5B). They include DN1s, DN2s and DN3s, and this split-GAL4 also targets all LNds in the lateral part of the brain; they control E activity. These data reinforce previous results ^23^ and indicate that the glutamatergic subset of DN1ps likely controls E activity and sleep at least in part through their interactions with other clock neurons. The upstream influence of dopamine on these cells and their functions widens the influence of the environment and brain state over the siesta and E-cell timing (see Discussion).

## Discussion

We show here that GPCRs are strongly expressed in the fly brain CCNN and can even define clock neuron identity. To identify individual receptors that contribute to specific neuron function, we combined a behavioral-sleep screen with a previously validated CRISPR-Cas9 specific neuron-mutagenesis strategy that exploited a new and comprehensive *Drosophila* GPCR guide library. The strategy was verified with a targeted sequencing approach and revealed a role of dopamine in sleep promotion. Dopamine generally inhibits sleep by stimulating locomotor activity, in flies as well as mammals ^35–37,41^. For example, compounds like amphetamine increase synaptic dopamine levels, which enhance fly activity and inhibit sleep ^42^. However, a specific sleep-promoting subpopulation of DN1ps uses this neurotransmitter for the opposite purpose, namely to promote sleep. This surprising conclusion resulted from the identification of the two dopamine receptors Dop1R1 and Dop1R2 as well as a clock neuron subpopulation, within which the two receptors gate the timing of daytime sleep.

The CCNN is an ideal platform to study neuromodulation, as genetic studies underscore the importance of neuropeptides to circadian behavior and even to circadian neuron subtype identification ^9,21,30^. This is also true for the mammalian brain and its CCNN, the suprachiasmatic nucleus (SCN) ^43^. Notably, all neuropeptides act through GPCRs, and 22 of the GPCRs expressed within the fly brain are at least 2x upregulated in the CCNN (Fig. 1E). This result is even more striking in light of our single cell RNAseq data, which show that many GPCRs are differentially expressed in clock neurons (Fig. 2).

To verify some of these expression patterns, we co-stained relevant GAL4 knock-in lines with an anti-PER antibody and thereby determined the overlap between receptor and clock protein gene expression. For example, the sequencing data suggest that *DH31-R* is highly enriched in the lLNv neurons and slightly enriched in the sLNv neurons and the DN1ps. The endogenous GAL4 line is strikingly consistent with this pattern: 3 out of 4 lLNvs were GFP-positive, whereas only 1 out of 4 sLNv neurons was labeled, explaining the expression difference between large and small LNvs. No LNds were labeled as predicted, but there was also an unexpected lack of DN1p expression in most brains. A *sNPFR* knock-in line was similarly consistent with the RNAseq data; it labeled DN1ps and only 1 out of 4 lLNvs (Fig. 2C). Similar differences between individual cells within a supposed uniform cluster were seen quite often, namely, with other GPCRs and other clock neuron subtypes (data not shown). These differences could reflect quantitative differences in expression between individual cells and low levels of GPCR mRNA expression as indicated by the single cell data, namely below threshold detection limits for immunostaining. The differences may also reflect stochastic differences between neighboring cells or unanticipated qualitative gene expression heterogeneity between cells, at the transcriptional, post-transcriptional or even translational level.

The data underscore more generally the importance of GPCR expression to the CCNN. To investigate this further, we analyzed our single cell data based only on GPCR expression. There were only marginal changes in cluster formation, i.e., we were still able to generate 17 clusters which were similar to those previously published. This surprising result indicates that GPCRs can define cellular identity at least within the clock system. They also likely define functional identity. This likely includes subtle contributions to phenotype beyond defining cell-specific ligand responses. For example, control flies usually have two very distinct subsets of DN1 neurons, some with low PER levels and others with higher PER levels. Removing dopamine receptors reduces this distinct separation and shows more neurons with a medium level of PER staining intensity (data not shown).

GPCRs are difficult to study genetically in mammals. This is because there are usually multiple genes encoding a GPCR. The situation is simpler in flies where there is usually only one gene that encodes each of the 124 GPCRs encoded in the fly genome. Our guide library targets all of these GPCRs and uses three independent guides for each receptor, a strategy shown to mutate a gene of interest in >95% of cases ^26–28^. As there are no available antibodies for GPCRs, we validated the knockouts by establishing a targeted genomic sequencing approach from isolated cells. The advantage of this approach is that we could directly investigate the effects of the guide-mediated mutagenesis in the clock network. This allowed us to directly interrogate their functionality by mutating these cells. Like the guide strategy previously used to eliminate PER ^27,28^, the GPCR guides can reliably delete GPCR-encoding genomic DNA from GFP-positive cells with no detectable deletions in GFP-negative cells. This further supports the notion that the UAS constructs suffer minimally if at all from background mutations (Fig. 3).

We focused on DN1ps because of their molecular complexity as well as their known contributions to M activity, the siesta and even nighttime sleep ^44^. Of the 21 DN1p-enriched GPCRs, 4 promote sleep based on the knockdown results. One of them, *rh7*, reproduced previously published results of whole body mutations, suggesting that it contributes to the siesta at least in part via DN1p expression ^34^. To our surprise, two dopamine receptors, *Dop1R1* and *Dop1R2*, also promote siesta sleep via the DN1ps and do so by gating the timing of siesta termination. This role resembles previous results indicating that *Vglut* and *AstC* are expressed in the DN1 neurons and influence the siesta and/or E activity under conditions that mimic seasonal regulation ^8,23,40^.

There are 4 different clusters of DN1 neurons that show elevated expression of *Dop1R1* or *Dop1R2*, suggesting that 1 or more of these clusters are responsible for the siesta phenotype. Unfortunately, the lack of more narrow drivers precludes more precise identification.

How can dopaminergic input influence the circadian gating of siesta sleep? The trans-tango pattern correlates nicely with previously published imaging data: lateral as well as dorsal clock neurons increase their cAMP levels in response to bath-applied dopamine ^45^. Bath application of the neuropeptide PDF causes a similar cAMP increase ^46^. Notably, this increase stabilizes PER, which is thought to delay the timing of the molecular clock ^47^. A similar mechanism might apply to dopamine and the DN1ps, which would then delay clock timing within these cells. Removing dopamine receptors would then lead to a decrease in cAMP levels and a consequent advance in timing, thereby explaining the early termination of the siesta in our experiments. Independent of mechanistic speculation, our data add to our view of how the CCNN works: dopaminergic input presumably reflects the monitoring by the CCNN of brain and environmental status, which then adjusts circadian timing before sending out time of day information to the rest of the animal (Fig. 6).

**Figure 6:**
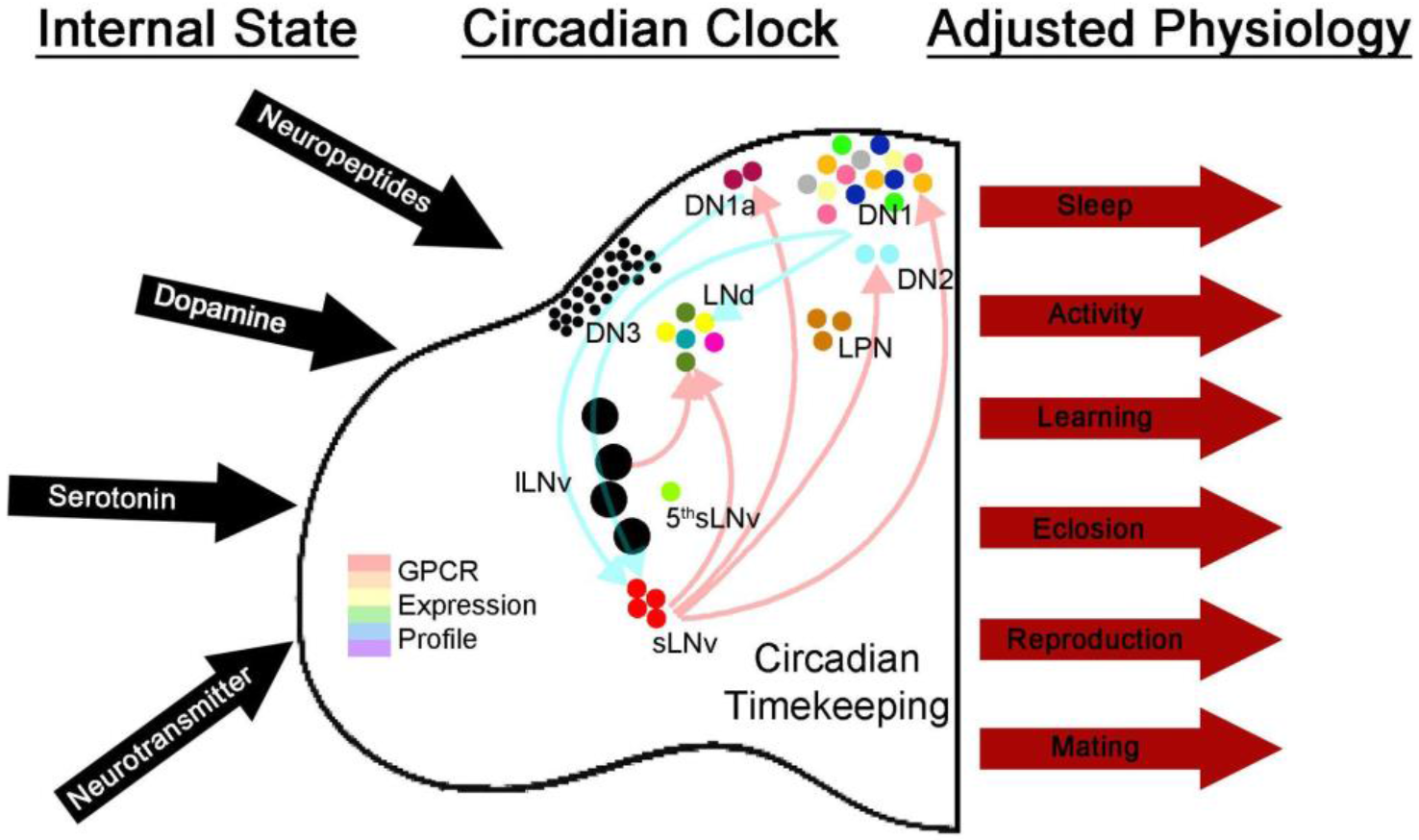
GPCR diversity contributes to the assessment and adaptation of animal physiology. The clock neuron network expresses approximately ⅔ of all GPCRs encoded by the fly genome. This allows the network to constantly assess the outside environment as well as the state of other networks in the brain which can then signal to the clock center. These signals can then alter the physiology of the clock and lead to changes in clock-controlled output.

Will the mutagenesis strategy and guide library be useful beyond the CCNN? We have found them to be failsafe, essentially 100% effective, far superior to current RNAi strategies and with no background issues. Given the broad role of neuropeptides, transmitters and GPCRs in most aspects of brain function and behavior, we anticipate that this GPCR mutagenesis strategy will be of use to a broad range of fly brain neuroscientists, well beyond the circadian system and the few other researchers who have already used them ^48^.

## Material and Methods

### Fly strains and rearing

Flies used in this study are listed in Table S1. All flies were raised at 25°C in a temperature-controlled incubator in LD 12:12.

### Generation of new fly lines

To generate UAS-guideRNA flies, we used the pCFD6 vector (addgene #73915, described in ^26^). In short, we generated three guides targeting the coding sequence of each GPCR. To identify possible target sites and avoid off-target effects, we used the optimal target finder developed by C. Dustin Rubinstein, Ed O’Connor-Giles & Kate M. O’Connor-Giles ^49^. Gene-specific guide sequences were then incorporated into the primers (see Suppl. Table 2) and the protocol described in ^26^ was followed. Correct clones were identified by colony PCR followed by sanger sequencing. Plasmids were injected into the attP1 site on the second chromosome (BDSC: 8621) by Rainbow Transgenic (Rainbow transgenic flies Inc., Camarillo, CA, USA). Individual flies were crossed to *w*^*1118*^ (BDSC: 3605) and screened for red eye color. Red eyed flies were balanced using *w;CyO/Sco;MKRS/TM6B* (BDSC: 3703).

### FACS sorting of *Drosophila* neurons

Neurons of interest were genetically labelled with *eGFP* using *nSyb*-GAL4 (for all neurons) or *clk856*-GAL4 (for clock neurons). About 2 week old male flies were entrained to the appropriate light:dark cycles. Brains of flies of the appropriate genotypes were dissected at ZT02 and ZT14 in ice cold Schneider’s medium (SM) and stored on ice until all dissections were completed (about 10 for *nSyb*, and 30 for *clk856*). Brains were incubated with enzyme solution (SM with 0.75ug/ul collagenase and 0.04 ug/ul dispase) at RT for 30 mins and then washed with SM, and triturated in SM. The volume was then brought up to ∼1ml and filtered through a 40 micron filter fitted on a FACS tube. GFP+ neurons of interest were isolated using the FACS Melody (BD) with sorting gates set by comparing to a GFP-negative neuron sample. 500 clock neurons or 1000 nSyb neurons were collected per sample in 100ul lysis buffer (Dynabeads mRNA direct kit) and frozen on dry ice immediately after collection. Two sets of neurons were collected from each dissociated sample.

### cDNA synthesis and library preparation for bulk sequencing

PolyA mRNA was isolated from the frozen cell samples using the Dynabeads mRNA direct kit (Thermo fisher 61011). Subsequently, cDNA was prepared using the method described in Picelli et. al. 2014 ^50^. cDNA integrity and concentration were assessed using a High Sensitivity D5000 ScreenTape (Agilent 5067-5592). ∼500pg of cDNA was used as input to make sequencing libraries with the Illumina Nextera XT DNA Library Preparation Kit (FC-131-1096) with 9 PCR cycles. Final libraries were quantified on a High Sensitivity D1000 ScreenTape on the TapeStation (5067-5584).

Libraries were run on the Illumina Nextseq 550 sequencing system. Reads were aligned to the dm6 version of the *Drosophila* genome using STAR ^51^. PCR duplicates were removed using Picard Tools (Picard Toolkit 2019. Broad Institute, GitHub Repository. https://broadinstitute.github.io/picard/; Broad Institute). Differential expression analysis between *nSyb* neurons and clock neurons was performed using the Bioconductor package edgeR ^52^.

### Single-cell RNA sequencing analysis

For details on single-cell RNAseq procedures refer to Ma et al. 2021 ^25^. In order to identify differentially expressed GPCRs in clock neurons, we first computed all marker genes in each cluster using the *FindAllMarkers* function of the Seurat package. Using a negative binomial generalized linear model, the batch effect from sequencing depth and conditions was regressed out. We next used an adjusted *p*-value significance of 0.05 and fold change cutoff of 1.25. GPCRs matching these criteria are regarded as differentially expressed in clock neurons. Their expression was plotted by the ComplexHeatmap package.

Annotated single-cell clustering data was used as the basis for GPCR-based reclustering using Seurat V4 in R^53^. For the downstream analysis, only cells from the 17 high-confidence annotated clusters were used. GPCR-based reclustering was done by restricting the variable features to GPCRs by setting the ‘features’ argument of the ‘ScaleData’ Seurat function to all genes identified as GPCRs from Hanlon and Andrew, 2015^4^ that were detectable in all timepoints. For the standard clustering, the FindVariableFeatures function with default settings was used to determine variable genes.

For both approaches, principal components (PCs) were determined using the RunPCA function, and the first 20 PCs were selected for clustering based on visual inspection of the ElbowPlots. Communities were generated using the standard workflow functions FindNeighbors, FindClusters, and RunUMAP with default settings. The ‘resolution’ argument of the FindClusters argument was set at the default of 0.8 after experimentation with resolutions as low as 0.5. Higher values for resolution and increased numbers of PCs did not improve clustering results separately or in combination for the GPCR clustering.

The cell cluster identities published in Ma, *et al*. ^25^ were stored in the Seurat metadata and used to assess the correspondence between the annotated clusters and the GPCR-based clustering. In general, the GPCR clusters correspond to a single annotated cluster and vice versa. There are two exceptions: the fusion of the two LNd clusters 9 and 12 into a single cluster and the fission of the DN1p cluster 4 into two similarly-sized clusters. The split of the DN1p cluster is characterized by a nearly twofold difference in average levels of *FMRFaR*, which is expressed in subsets of DN1p neurons.

### Targeted genomic sequencing

To analyze the potency of our guide library we established a targeted genomic sequencing approach similar to the 16S metagenomic sequencing library preparation protocol (Illumina #15044223). In short, we generated 3 pairs of primers for each gene (Table S3). Each pair of primers was designed to amplify 230-270 bp long genomic regions centered around the predicted guide cut sites. Each primer pair was tested using *w*^*1118*^ genomic DNA and yielded the desired bands only (data not shown). To analyze the potency of our guide library to mutate genes in a cell-specific manner, we labeled the clock network with GFP (*clk856>EGFP*) and expressed *Cas9* along with the guide of interest. We FACS sorted (see above) 2000 GFP+ and 2000 GFP-cells and extracted DNA using a DNA extraction buffer. We then used this extract and performed a PCR with all six primers for a gene of interest using ExTaq polymerase. After Dynabeads mediated cleanup, adapters were ligated followed by another round of cleanup. Final libraries were quantified on a High Sensitivity D1000 ScreenTape on the TapeStation (5067-5584). Libraries were then pooled and sequenced using the miseq platform (Genewiz Inc.). Libraries were analyzed using Crispresso2 ^54^.

### Behavioral analysis

2-7 days old male flies were individually placed into glass tubes with food (2% agar, 4% sucrose) on one end and a plug to close the tube on the other end. These tubes were subsequently placed into *Drosophila* Activity Monitors (DAM, Trikinetics Inc, Waltham, MA, USA) and a computer recorded the number of IR light beam interruptions in one minute intervals. Flies were recorded for one week in LD 12:12 followed by DD for another week. Each experiment was performed at least twice.

To allow for proper entrainment to the LD cycle, we used the last 4 days of LD to analyze sleep; defined as 5 minutes of inactivity ^55,56^. We then created average sleep profiles by averaging the amount of sleep across days and flies for each genotype in half hour bins. To analyze sleep in more detail, we split the sleep amount into 4 sections of 6 hours each: Morning (ZT21-ZT3), Siesta (ZT3-ZT9), Evening (ZT9-ZT15) and Night (ZT15-ZT21). Individual values were plotted as scatter plots and statistical analysis was performed using a one-way ANOVA followed by a post-hoc Tukey test (astatsa.com). p<0.05 compared to all control groups was considered significant.

To analyze changes in rhythmicity we performed a chi-square analysis for rhythmicity and analyzed the speed of the clock. Individual values were compared using a one-way ANOVA followed by a post-hoc Tukey test (astatsa.com) p<0.05 compared to all control groups was considered statistically significant.

### Immunohistochemistry

2-7 days old male flies were fixed infor 2h 45min in 4% PFA in PBST (phosphate buffered saline including 0.5% TritonX). After rinsing 5×10min each in PBST, brains were dissected and blocked for 2h in 5% normal goat serum (NGS) in PBST. Primary antibodies were applied overnight at room temperature (RT). Primary antibodies were the following: chicken anti-GFP (1:1500, abcam), rabbit anti-PER (1:1000 ^57^), mouse anti-PDF (1:1000, DSHB) and rat anti-RFP (1:500, chromotec). After rinsing 5x with PBST, secondary antibodies (1:200, Alexa Fluor, Fisher Scientific) were applied for 2h at RT followed by rinsing 5×10min each with PBST. Brains were subsequently mounted on glass slides using Vectashield mounting medium (Vector laboratories) and imaged using the Leica SP5 confocal microscope. For PER quantification we used a 3×3 pixel area to determine the staining intensity of neurons and corrected by measuring 3 different background intensities as previously described ^58^.

## Acknowledgements

We want to thank Dr. R. Stanewsky, Dr, Y Rao, Dr. R. Allada and Dr. C. Helfrich-Förster for their fly lines and antibodies. A special thank you to Dr. K. Clement for the help with setting up Crispresso2 on our server. The PDF antibody developed by Dr. J. Blau was obtained from the Developmental Studies Hybridoma Bank, created by the NICHD of the NIH and maintained at The University of Iowa, Department of Biology, Iowa City, IA 52242. The pCFD6 plasmid was generated by Dr. F. Port and Dr. S. Bullock. Stocks obtained from the Bloomington *Drosophila* Stock Center (NIH P40OD018537) were used in this study. M.S. was funded by a Research Fellowship of the German Research Foundation (DFG). This study was further funded by the Howard Hughes Medical Institute.

## References

1. Griffith, L. C. Neuromodulatory control of sleep in Drosophila melanogaster: integration of competing and complementary behaviors. Curr. Opin. Neurobiol. 23, 819–823 (2013).

2. Marder, E. Neuromodulation of Neuronal Circuits: Back to the Future. Neuron vol. 76 1–11 (2012).

3. Yang, D. et al. G protein-coupled receptors: structure-and function-based drug discovery. Signal Transduct Target Ther 6, 7 (2021).

4. Hanlon, C. D. & Andrew, D. J. Outside-in signaling--a brief review of GPCR signaling with a focus on the Drosophila GPCR family. J. Cell Sci. 128, 3533–3542 (2015).

5. Renn, S. C., Park, J. H., Rosbash, M., Hall, J. C. & Taghert, P. H. A pdf neuropeptide gene mutation and ablation of PDF neurons each cause severe abnormalities of behavioral circadian rhythms in Drosophila. Cell 99, 791–802 (1999).

6. Hermann-Luibl, C., Yoshii, T., Senthilan, P. R., Dircksen, H. & Helfrich-Förster, C. The ion transport peptide is a new functional clock neuropeptide in the fruit fly Drosophila melanogaster. J. Neurosci. 34, 9522–9536 (2014).

7. Fujiwara, Y. et al. The CCHamide1 Neuropeptide Expressed in the Anterior Dorsal Neuron 1 Conveys a Circadian Signal to the Ventral Lateral Neurons in. Front. Physiol. 9, 1276 (2018).

8. Díaz, M. M., Schlichting, M., Abruzzi, K. C., Long, X. & Rosbash, M. Allatostatin-C/AstC-R2 Is a Novel Pathway to Modulate the Circadian Activity Pattern in Drosophila. Curr. Biol. 29, 13–22.e3 (2019).

9. Yao, Z. & Shafer, O. T. The Drosophila circadian clock is a variably coupled network of multiple peptidergic units. Science 343, 1516–1520 (2014).

10. Ahmad, M., Li, W. & Top, D. Integration of Circadian Clock Information in the Drosophila Circadian Neuronal Network. Journal of Biological Rhythms vol. 36 203–220 (2021).

11. Helfrich-Förster, C. The period clock gene is expressed in central nervous system neurons which also produce a neuropeptide that reveals the projections of circadian pacemaker cells within the brain of Drosophila melanogaster. Proc. Natl. Acad. Sci. U. S. A. 92, 612–616 (1995).

12. Grima, B., Chélot, E., Xia, R. & Rouyer, F. Morning and evening peaks of activity rely on different clock neurons of the Drosophila brain. Nature 431, 869–873 (2004).

13. Stoleru, D., Peng, Y., Agosto, J. & Rosbash, M. Coupled oscillators control morning and evening locomotor behaviour of Drosophila. Nature 431, 862–868 (2004).

14. Shang, Y., Griffith, L. C. & Rosbash, M. Light-arousal and circadian photoreception circuits intersect at the large PDF cells of the Drosophila brain. Proc. Natl. Acad. Sci. U. S. A. 105, 19587–19594 (2008).

15. Sheeba, V. et al. Large ventral lateral neurons modulate arousal and sleep in Drosophila. Curr. Biol. 18, 1537–1545 (2008).

16. Yoshii, T. et al. The neuropeptide pigment-dispersing factor adjusts period and phase of Drosophila’s clock. J. Neurosci. 29, 2597–2610 (2009).

17. Kaneko, H. et al. Circadian rhythm of temperature preference and its neural control in Drosophila. Curr. Biol. 22, 1851–1857 (2012).

18. Veleri, S., Brandes, C., Helfrich-Förster, C., Hall, J. C. & Stanewsky, R. A self-sustaining, light-entrainable circadian oscillator in the Drosophila brain. Curr. Biol. 13, 1758–1767 (2003).

19. Nettnin, E. A., Sallese, T. R., Nasseri, A., Saurabh, S. & Cavanaugh, D. J. Dorsal clock neurons in sculpt locomotor outputs but are dispensable for circadian activity rhythms. iScience 24, 103001 (2021).

20. Klarsfeld, A. Novel Features of Cryptochrome-Mediated Photoreception in the Brain Circadian Clock of Drosophila. Journal of Neuroscience vol. 24 1468–1477 (2004).

21. Yao, Z., Bennett, A. J., Clem, J. L. & Shafer, O. T. The Drosophila Clock Neuron Network Features Diverse Coupling Modes and Requires Network-wide Coherence for Robust Circadian Rhythms. Cell Rep. 17, 2873–2881 (2016).

22. Guo, F., Holla, M., Díaz, M. M. & Rosbash, M. A Circadian Output Circuit Controls Sleep-Wake Arousal in Drosophila. Neuron 100, 624–635.e4 (2018).

23. Guo, F. et al. Circadian neuron feedback controls the Drosophila sleep--activity profile. Nature 536, 292–297 (2016).

24. Lamaze, A., Krätschmer, P., Chen, K.-F., Lowe, S. & Jepson, J. E. C. A Wake-Promoting Circadian Output Circuit in Drosophila. Curr. Biol. 28, 3098–3105.e3 (2018).

25. Ma, D. et al. A transcriptomic taxonomy of circadian neurons around the clock. Elife 10, (2021).

26. Port, F. & Bullock, S. L. Augmenting CRISPR applications in Drosophila with tRNA-flanked sgRNAs. Nat. Methods 13, 852–854 (2016).

27. Delventhal, R. et al. Dissection of central clock function in through cell-specific CRISPR-mediated clock gene disruption. Elife 8, (2019).

28. Schlichting, M., Díaz, M. M., Xin, J. & Rosbash, M. Neuron-specific knockouts indicate the importance of network communication to Drosophila rhythmicity. eLife vol. 8 (2019).

29. Bulthuis, N., Spontak, K. R., Kleeman, B. & Cavanaugh, D. J. Neuronal Activity in Non-LNv Clock Cells Is Required to Produce Free-Running Rest:Activity Rhythms in Drosophila. J. Biol. Rhythms 34, 249–271 (2019).

30. Abruzzi, K. C. et al. RNA-seq analysis of Drosophila clock and non-clock neurons reveals neuron-specific cycling and novel candidate neuropeptides. PLoS Genet. 13, e1006613 (2017).

31. Deng, B. et al. Chemoconnectomics: Mapping Chemical Transmission in Drosophila. Neuron 101, 876–893.e4 (2019).

32. Hyun, S. et al. Drosophila GPCR Han Is a Receptor for the Circadian Clock Neuropeptide PDF. Neuron vol. 48 267–278 (2005).

33. Schlichting, M. et al. Light-Mediated Circuit Switching in the Drosophila Neuronal Clock Network. Curr. Biol. 29, 3266–3276.e3 (2019).

34. Kistenpfennig, C. et al. A New Rhodopsin Influences Light-dependent Daily Activity Patterns of Fruit Flies. J. Biol. Rhythms 32, 406–422 (2017).

35. Kume, K. Dopamine Is a Regulator of Arousal in the Fruit Fly. Journal of Neuroscience vol. 25 7377–7384 (2005).

36. Wu, M. N., Koh, K., Yue, Z., Joiner, W. J. & Sehgal, A. A genetic screen for sleep and circadian mutants reveals mechanisms underlying regulation of sleep in Drosophila. Sleep 31, 465–472 (2008).

37. Riemensperger, T. et al. Behavioral consequences of dopamine deficiency in the Drosophila central nervous system. Proceedings of the National Academy of Sciences vol. 108 834–839 (2011).

38. Pimentel, D. et al. Operation of a homeostatic sleep switch. Nature 536, 333–337 (2016).

39. Konopka, R. J. & Benzer, S. Clock mutants of Drosophila melanogaster. Proc. Natl. Acad. Sci. U. S. A. 68, 2112–2116 (1971).

40. Chatterjee, A. et al. Reconfiguration of a Multi-oscillator Network by Light in the Drosophila Circadian Clock. Curr. Biol. 28, 2007–2017.e4 (2018).

41. Qu, W.-M. et al. Essential role of dopamine D2 receptor in the maintenance of wakefulness, but not in homeostatic regulation of sleep, in mice. J. Neurosci. 30, 4382–4389 (2010).

42. Karam, C. S., Williams, B. L., Jones, S. K. & Javitch, J. A. The Role of the Dopamine Transporter in the Effects of Amphetamine on Sleep and Sleep Architecture in Drosophila. Neurochem. Res. (2021) doi:10.1007/s11064-021-03275-4.

43. Ono, D., Honma, K.-I. & Honma, S. Roles of Neuropeptides, VIP and AVP, in the Mammalian Central Circadian Clock. Front. Neurosci. 15, 650154 (2021).

44. Lamaze, A. & Stanewsky, R. DN1p or the ‘Fluffy’ Cerberus of Clock Outputs. Front. Physiol. 10, 1540 (2019).

45. Fernandez-Chiappe, F. et al. Dopamine signaling in wake promoting clock neurons is not required for the normal regulation of sleep in Drosophila. doi:10.1101/2020.05.20.106369.

46. Shafer, O. T. et al. Widespread receptivity to neuropeptide PDF throughout the neuronal circadian clock network of Drosophila revealed by real-time cyclic AMP imaging. Neuron 58, 223–237 (2008).

47. Li, Y., Guo, F., Shen, J. & Rosbash, M. PDF and cAMP enhance PER stability in Drosophila clock neurons. Proc. Natl. Acad. Sci. U. S. A. 111, E1284–90 (2014).

48. Vogt, K. et al. Internal state configures olfactory behavior and early sensory processing in Drosophila larvae. doi:10.1101/2020.03.02.973941.

49. Gratz, S. J. et al. Highly specific and efficient CRISPR/Cas9-catalyzed homology-directed repair in Drosophila. Genetics 196, 961–971 (2014).

50. Picelli, S. et al. Full-length RNA-seq from single cells using Smart-seq2. Nat. Protoc. 9, 171–181 (2014).

51. Dobin, A. et al. STAR: ultrafast universal RNA-seq aligner. Bioinformatics 29, 15–21 (2013).

52. Robinson, M. D., McCarthy, D. J. & Smyth, G. K. edgeR: a Bioconductor package for differential expression analysis of digital gene expression data. Bioinformatics 26, 139–140 (2010).

53. Hao, Y. et al. Integrated analysis of multimodal single-cell data. Cell 184, 3573–3587.e29 (2021).

54. Clement, K. et al. CRISPResso2 provides accurate and rapid genome editing sequence analysis. Nat. Biotechnol. 37, 224–226 (2019).

55. Hendricks, J. C. et al. Rest in Drosophila Is a Sleep-like State. Neuron vol. 25 129–138 (2000).

56. Shaw, P. J. Correlates of Sleep and Waking in Drosophila melanogaster. Science vol. 287 1834–1837 (2000).

57. Stanewsky, R. et al. The cryb mutation identifies cryptochrome as a circadian photoreceptor in Drosophila. Cell 95, 681–692 (1998).

58. Menegazzi, P. et al. Drosophila clock neurons under natural conditions. J. Biol. Rhythms 28, 3–14 (2013).

