## Supplemental data for "Dopamine and GPCR-mediated modulation of DN1 clock neurons gates the circadian timing of sleep"

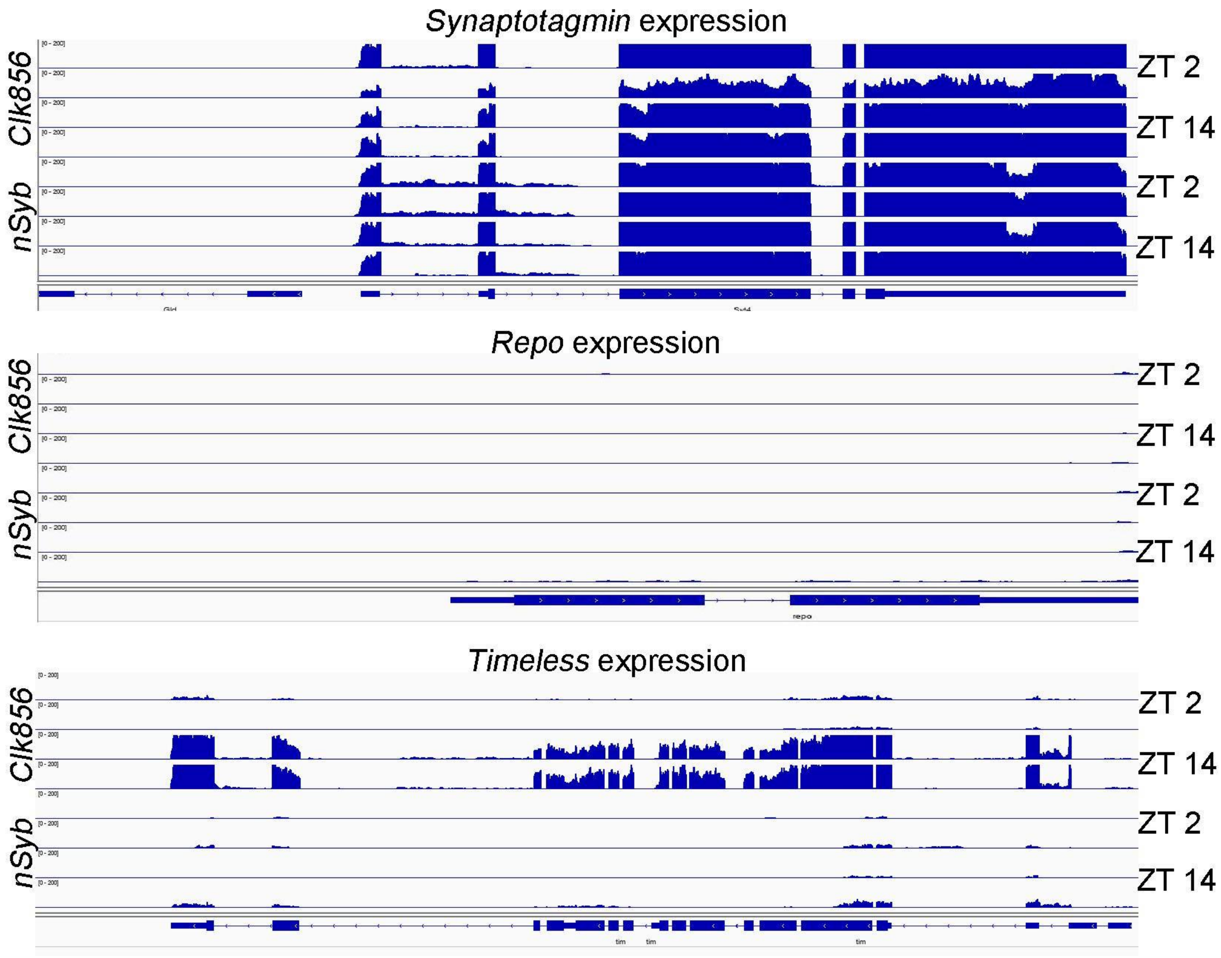


**Figure S1: Bulk sequencing of FACS sorted neurons shows expected enrichment of key genes.** Upper panel: Sorted clock neurons (*clk856>EGFP*) and sorted random neurons (*nSyb>EGFP*) show high expression of the neuronal marker *synaptotagmin*. Middle panel: Sorted clock neurons (*clk856>EGFP*) and sorted random neurons (*nSyb>EGFP*) show no expression of the glial marker *repo* indicating no contamination by glial cells. Lower Panel: Sorted clock neurons (*clk856>EGFP*) show high *timeless* expression at ZT14 and little expression at ZT2. Random neurons (*nSyb>EGFP*) show strongly reduced levels of *timeless* at all time points indicating successful enrichment of clock neurons. Maximum set to 200 reads for all graphs.

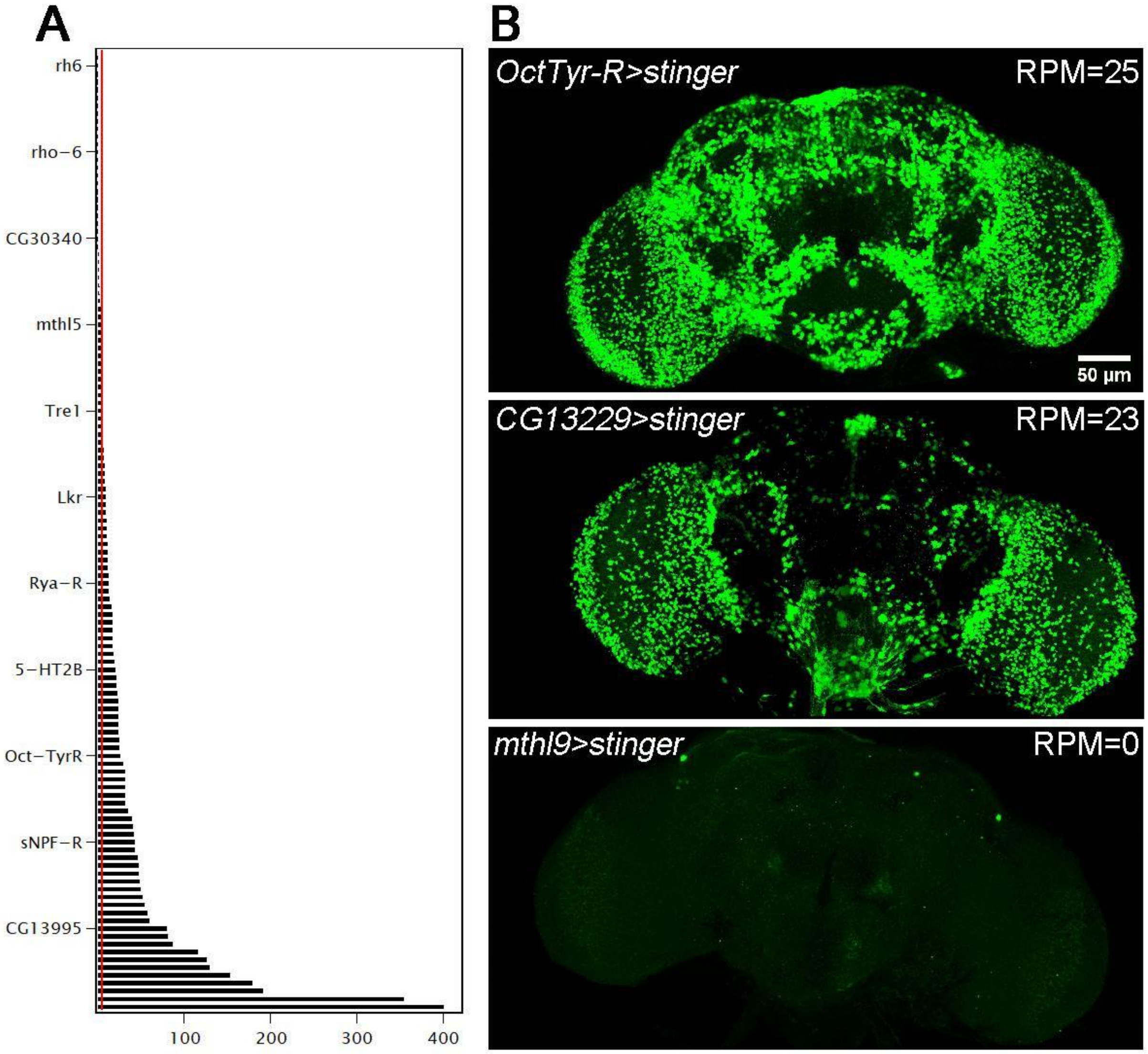


**Figure S2: GPCRs are differentially expressed in the brain.** **A** Normalized expression (RPM) of GPCRs from isolated neurons (*nSyb>GFP*). Expression of GPCRs varies strongly from gene to gene. **B** Whole mount brains of flies expressing nuclear GFP under the control of specific GPCRs. From top to bottom: *Oct-Tyr-R>stinger*, *CG13229>stinger* and *mthl9>stinger*. The number of nuclei labeled with GFP corresponds to the number of normalized reads as indicated in the top right corner of each image.

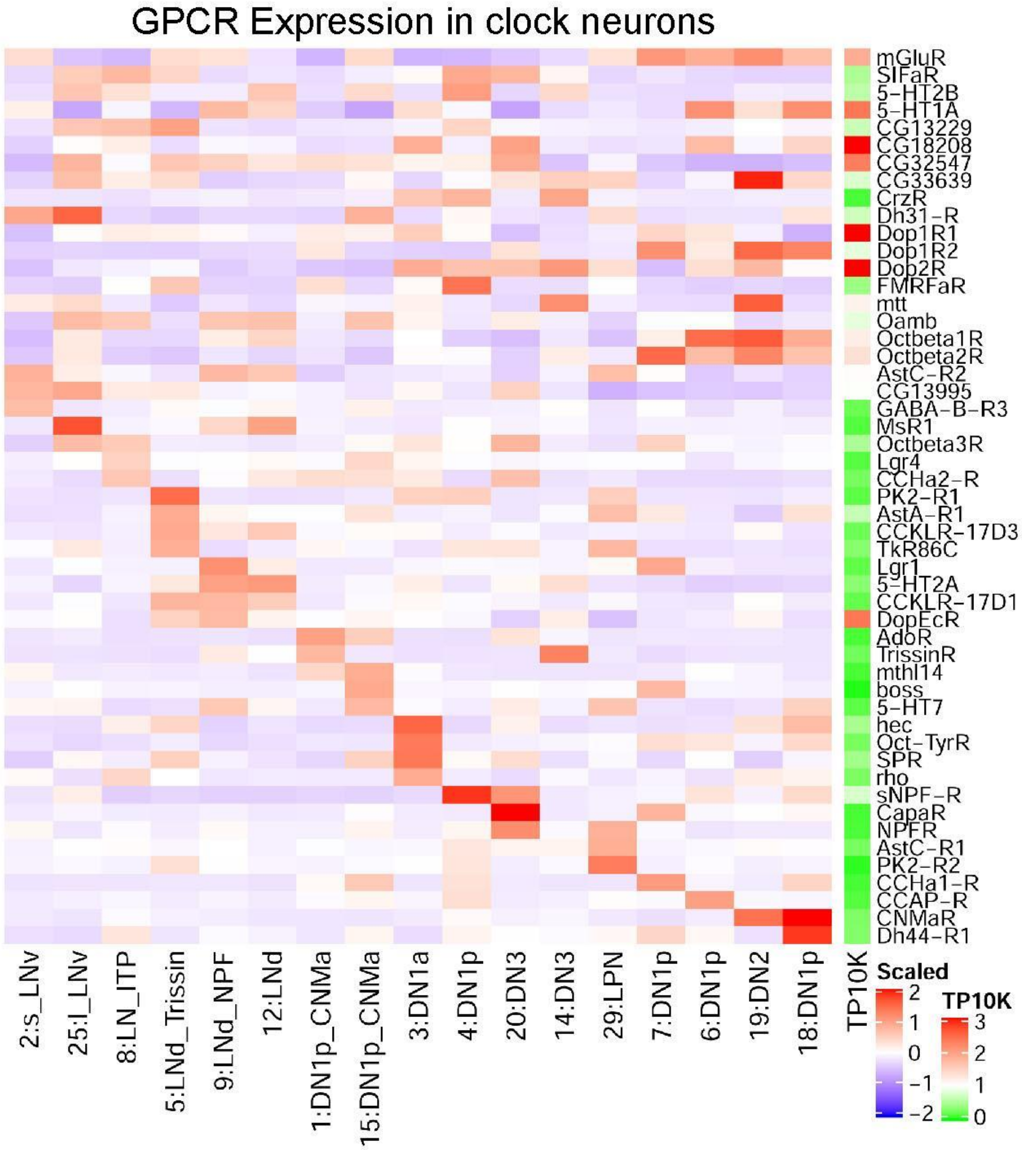


**Figure S3: GPCRs are highly differentially expressed in the clock neuron network based on single-cell RNA sequencing.** Heatmap on the right (green: low expression red: high expression) indicates the overall expression level of the specific GPCR. Heatmap on the left (blue: low expression red: high expression) represents the expression levels of specific GPCRs within the 17 high confidence clock neuron clusters.

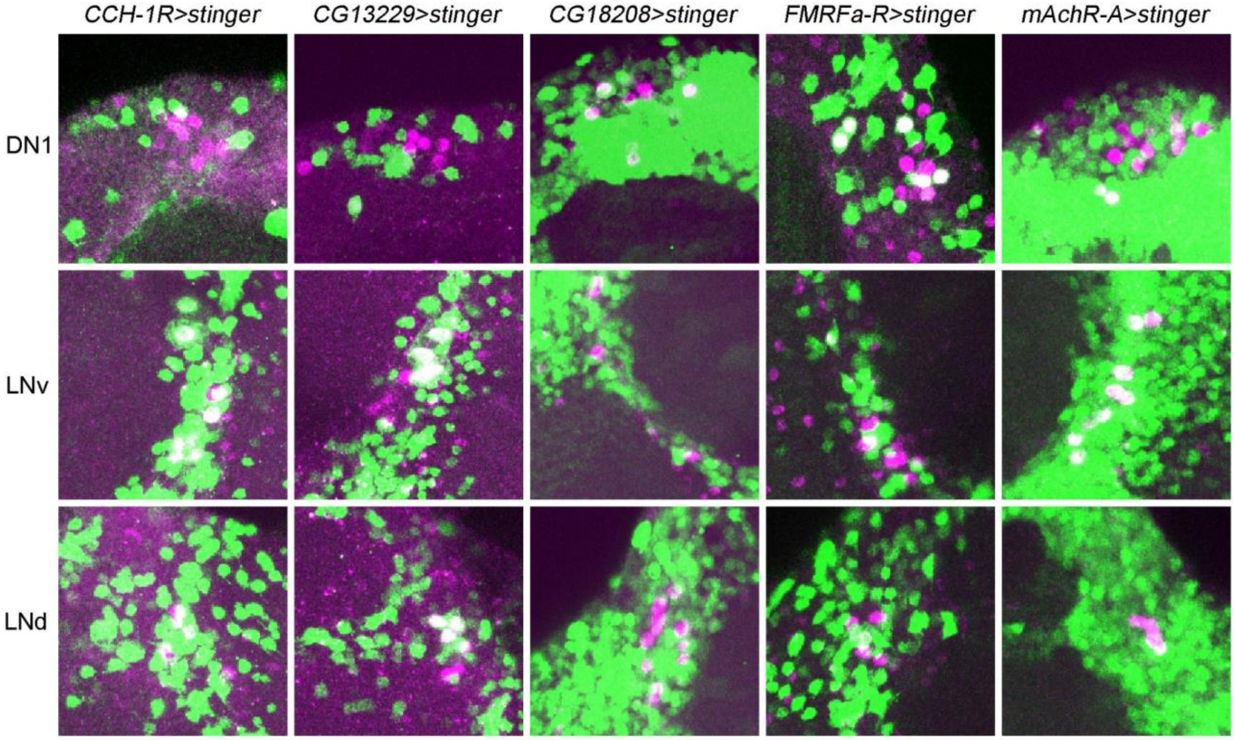


**Figure S4: GPCRs are differentially expressed based on endogenous GAL4 expression.** Whole mount brains were labeled with anti-GFP (green) and anti-PER (magenta). To analyze colocalization, we generated z-stacks of all the stacks containing the cells of interest. From top to bottom: DN1ps (posterior brain), ventro-lateral neurons (PDF-cells) and LNds. Genotypes are indicated on top of each row.

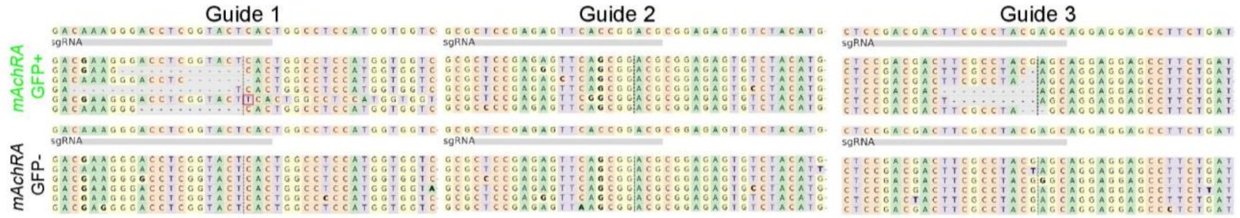


**Figure S5: Guide library efficiently mutates GPCRs of choice in a cell-type specific manner.** Mutagenesis of *mAchR-A* in clock neurons. Guides 1 and 3 cause deletions in GFP positive neurons, whereas no deletions were detected in Guide 2 and GFP negative cells.

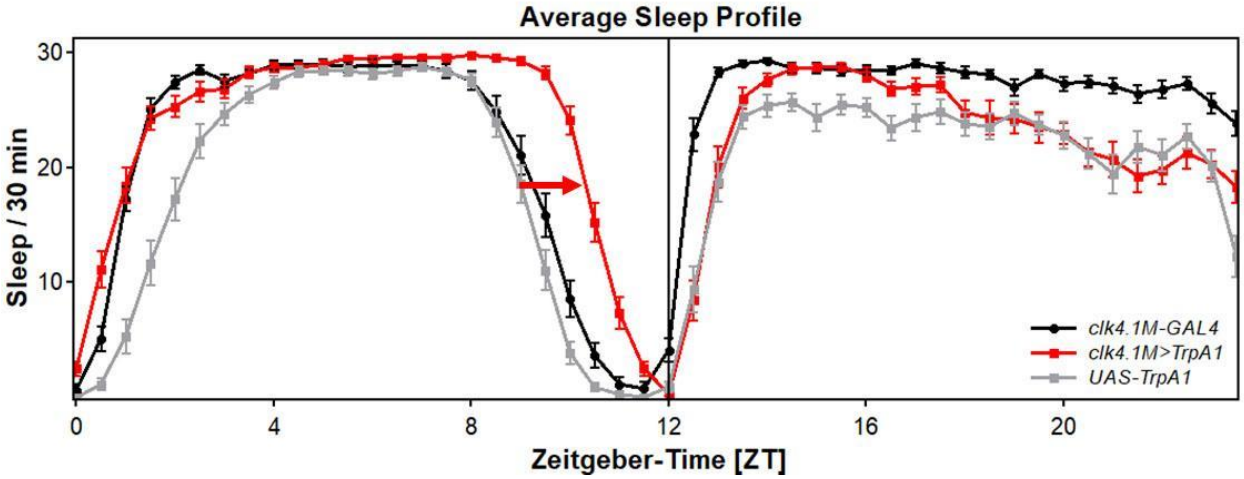


**Figure S6: Activating DN1 neurons increases the siesta.** Mild activation (26°C) of DN1ps using the temperature sensitive cation channel *TrpA1* increases sleep in the second half of the day (red line) compared to both controls (grey and black lines), whereas there were no major effects on sleep at any other time of the day.

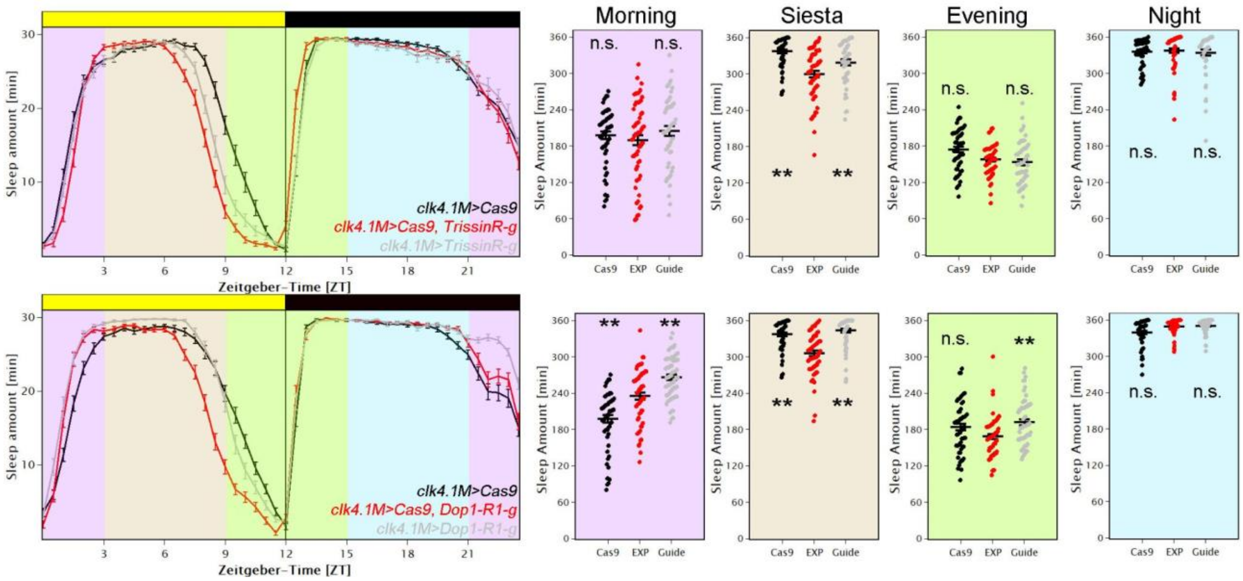


**Figure S7: Mutagenesis of *Dop1R1* and *TrissinR* in DN1ps affects siesta sleep** Upper panel: Average sleep profile of male flies in which *TrissinR* was mutated in the DN1ps (red) and controls (grey and black). Error bars represent SEM. Background colors indicate 4x6h periods which were quantified. Quantification of sleep is separated into 4 different time zones: Morning (ZT21-ZT3), Siesta (ZT3-ZT9), Evening (ZT9-ZT15) and Night (ZT15-ZT21). Removing *TrissinR* in the DN1ps significantly reduced sleep during the siesta ($p<0.01$), whereas other times of day were not affected. Lower panel: Average sleep profile of male flies in which *Dop1R1* was mutated in the DN1ps (red) and controls (grey and black). Error bars represent SEM. Background colors indicate 4x6h periods which were quantified. Quantification of sleep is separated into 4 different time zones: Morning (ZT21-ZT3), Siesta (ZT3-ZT9), Evening (ZT9-ZT15) and Night (ZT15-ZT21). Removing *Dop1R1* in the DN1ps significantly reduced sleep during the siesta ($p<0.01$), whereas other times of day were not affected.

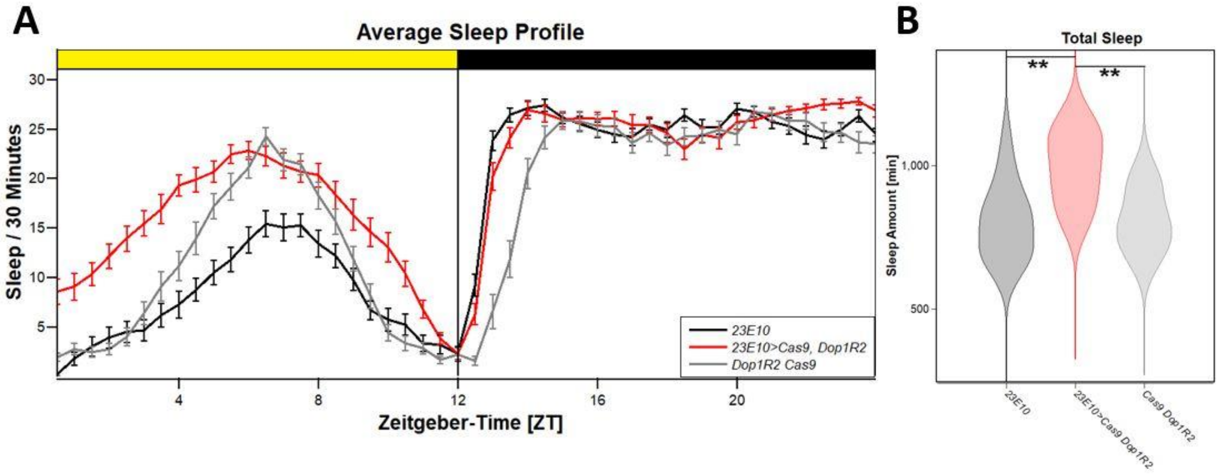


**Figure S8: Dop1R2 reduces sleep in the dorsal fan shaped body. A** Average sleep profile of female flies with mutated *Dop1R2* in the dFb (red) and controls (grey and black). Error bars represent SEM. Experimental flies show increased sleep especially during daytime. **B** Quantification of total sleep. Experimental flies show significantly increased sleep amounts compared to both controls ($p<0.01$).

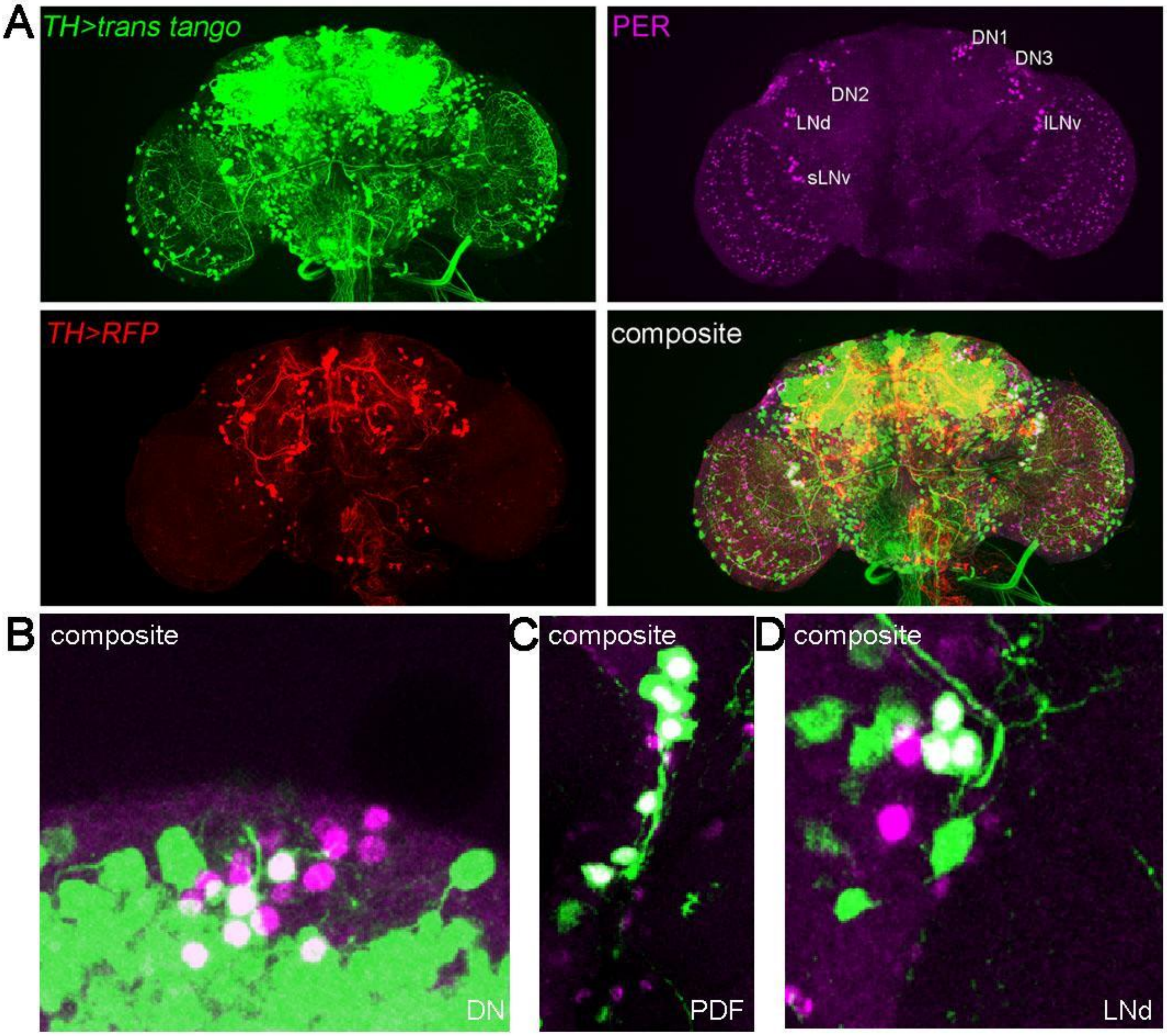


**Figure S9: Dopaminergic neurons connect to a variety of clock neurons. A** Whole mount brain of *TH>RFP, trans-tango; QUAS-Gcamp* stained with anti-GFP (green, labeling downstream targets of TH neurons), anti-RFP (red, labels TH neurons) and anti-PER (magenta, labeling clock cells). The TH neurons connect to many cells in the fly brain. **B** TH neurons connect to approximately 4 DN1ps and DN2s. **C** TH neurons connect to sLNvs and lLNvs. **D** TH neurons connect to 3 LNds.

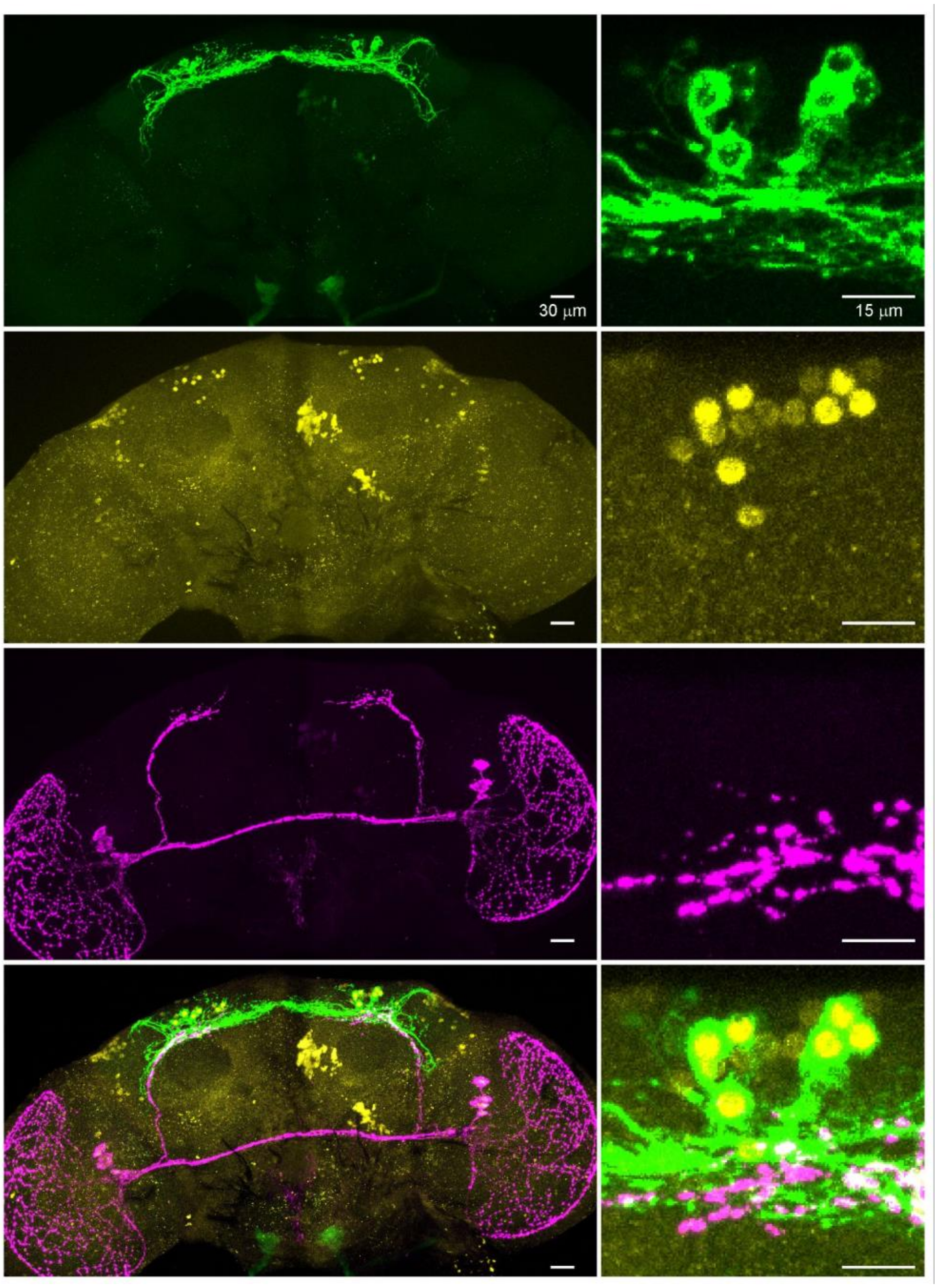


**Figure S10:** ***per-AD Vglut-DBD>EGFP* expressing flies have 7-8 GFP+ DN1ps.** Brains stained against anti-GFP (green), anti-PER (yellow) and anti-PDF (magenta).

Table S1: Flies used in this study

| Genotype | Description | Reference / Source |
|---|---|---|
| *clk856-GAL4* | Expresses GAL4 in most clock neurons | (Gummadova et al. 2009) |
| *UAS-EGFP* | Expresses EGFP under UAS control | BDSC 5430 |
| *nSyb-GAL4* | Expresses GAL4 in neurons | BDSC 51635 |
| *UAS-stinger* | Expresses nuclear GFP under UAS control | BDSC 84278 |
| *Dh31-R-Gal4-RA/C* | DKI0062 | (Deng et al. 2019) |
| *sNPF-R-Gal4* | WCKI1115 | (Deng et al. 2019) |
| *Oct-TyrR-GAL4-T1/Sb* | | (Deng et al. 2019) |
| *CG13229-Gal4* | DKI0133 | (Deng et al. 2019) |
| *mthl9-Gal4* | WCKI1036 | (Deng et al. 2019) |
| *FMRFaR-Gal4* | WCKI1117 | (Deng et al. 2019) |
| *CG18208-Gal4* | WCKI1100 | (Deng et al. 2019) |
| *CCHa1-R-Gal4* | DKI0077 | (Deng et al. 2019) |
| *mAChR-A-Gal4* | DKI0218 | (Deng et al. 2019) |
| *UAS-Cas9.P2 (attP40)* | Expresses Cas9 under UAS control | BDSC 58985 |
| *clk4.1M-GAL4* | Expresses GAL4 in DN1s | (Zhang et al. 2010) |
| *UAS-TrpA1* | | BDSC 26263 |
| *23E10-GAL4* | | BDSC 49032 |
| *per-AD* | | BDSC 70551 |
| *Vglut-DBD* | | BDSC 69036 |
| *UAS-trans-tango UAS-RFP QUAS-Gcamp* | | Modified from BDSC 77124 |
| *UAS-5-HT1A-g* | UAS-guide-RNA line | This study |
| *UAS-5HT1B-g* | UAS-guide-RNA line | This study |
| *UAS-5HT2A-g* | UAS-guide-RNA line | This study |
| *UAS-5HT2B-g* | UAS-guide-RNA line | This study |
| *UAS-5HT7-g* | UAS-guide-RNA line | This study |
| *UAS-AdoR-g* | UAS-guide-RNA line | This study |

| | | |
|---|---|---|
| *UAS-AkhR-g* | UAS-guide-RNA line | This study |
| *UAS-AstA-R1-g* | UAS-guide-RNA line | This study |
| *UAS-AstA-R2-g* | UAS-guide-RNA line | This study |
| *UAS-AstC-R1-g* | UAS-guide-RNA line | This study |
| *UAS-AstC-R2-g* | UAS-guide-RNA line | This study |
| *UAS-boss-g* | UAS-guide-RNA line | This study |
| *UAS-CapaR-g* | UAS-guide-RNA line | This study |
| *UAS-CCAP-R-g* | UAS-guide-RNA line | This study |
| *UAS-CCHa1-R-g* | UAS-guide-RNA line | This study |
| *UAS-CCHa2-R-g* | UAS-guide-RNA line | This study |
| *UAS-CCKLR-14D1-g* | UAS-guide-RNA line | This study |
| *UAS-CCKLR-17D3-g* | UAS-guide-RNA line | This study |
| *UAS-CG11318-g* | UAS-guide-RNA line | This study |
| *UAS-CG12290-g* | UAS-guide-RNA line | This study |
| *UAS-CG12796-g* | UAS-guide-RNA line | This study |
| *UAS-CG13229-g* | UAS-guide-RNA line | This study |
| *UAS-CG13575-g* | UAS-guide-RNA line | This study |
| *UAS-CG13579-g* | UAS-guide-RNA line | This study |
| *UAS-CG13995-g* | UAS-guide-RNA line | This study |
| *UAS-CG15556-g* | UAS-guide-RNA line | This study |
| *UAS-CG15614-g* | UAS-guide-RNA line | This study |
| *UAS-CG15744-g* | UAS-guide-RNA line | This study |
| *UAS-CG18208-g* | UAS-guide-RNA line | This study |
| *UAS-CG30340-g* | UAS-guide-RNA line | This study |
| *UAS-CG31760-g* | UAS-guide-RNA line | This study |
| *UAS-CG32447-g* | UAS-guide-RNA line | This study |
| *UAS-CG32547-g* | UAS-guide-RNA line | This study |
| *UAS-CG33639-g* | UAS-guide-RNA line | This study |
| *UAS-CG4313-g* | UAS-guide-RNA line | This study |
| *UAS-CG44153-g* | UAS-guide-RNA line | This study |
| *UAS-CG7497-g* | UAS-guide-RNA line | This study |
| *UAS-Cirl-g* | UAS-guide-RNA line | This study |

| | | |
|---|---|---|
| *UAS-CNMaR-g* | UAS-guide-RNA line | This study |
| *UAS-CrzR-g* | UAS-guide-RNA line | This study |
| *UAS-Dh31-R-g* | UAS-guide-RNA line | This study |
| *UAS-Dh44-R1-g* | UAS-guide-RNA line | This study |
| *UAS-Dh44-R2-g* | UAS-guide-RNA line | This study |
| *UAS-Dop1R1-g* | UAS-guide-RNA line | This study |
| *UAS-Dop1R2-g* | UAS-guide-RNA line | This study |
| *UAS-Dop2R-g* | UAS-guide-RNA line | This study |
| *UAS-DopEcR-g* | UAS-guide-RNA line | This study |
| *UAS-ETHR-g* | UAS-guide-RNA line | This study |
| *UAS-FMRFaR-g* | UAS-guide-RNA line | This study |
| *UAS-fz-g* | UAS-guide-RNA line | This study |
| *UAS-fz2-g* | UAS-guide-RNA line | This study |
| *UAS-fz3-g* | UAS-guide-RNA line | This study |
| *UAS-fz4-g* | UAS-guide-RNA line | This study |
| *UAS-GABA-B-R1-g* | UAS-guide-RNA line | This study |
| *UAS-GABA-B-R2-g* | UAS-guide-RNA line | This study |
| *UAS-GABA-B-R3-g* | UAS-guide-RNA line | This study |
| *UAS-hec-g* | UAS-guide-RNA line | This study |
| *UAS-Lgr1-g* | UAS-guide-RNA line | This study |
| *UAS-Lgr3-g* | UAS-guide-RNA line | This study |
| *UAS-Lgr4-g* | UAS-guide-RNA line | This study |
| *UAS-Lkr-g* | UAS-guide-RNA line | This study |
| *UAS-mAchR-A-g* | UAS-guide-RNA line | This study |
| *UAS-mAchR-B-g* | UAS-guide-RNA line | This study |
| *UAS-mAchR-C-g* | UAS-guide-RNA line | This study |
| *UAS-mGluR-g* | UAS-guide-RNA line | This study |
| *UAS-moody-g* | UAS-guide-RNA line | This study |
| *UAS-MsR1-g* | UAS-guide-RNA line | This study |
| *UAS-MsR2-g* | UAS-guide-RNA line | This study |
| *UAS-mth-g* | UAS-guide-RNA line | This study |
| *UAS-mthl1-g* | UAS-guide-RNA line | This study |

| | | |
|---|---|---|
| *UAS-mthl10-g* | UAS-guide-RNA line | This study |
| *UAS-mthl11-g* | UAS-guide-RNA line | This study |
| *UAS-mthl12-g* | UAS-guide-RNA line | This study |
| *UAS-mthl13-g* | UAS-guide-RNA line | This study |
| *UAS-mthl14-g* | UAS-guide-RNA line | This study |
| *UAS-mthl15-g* | UAS-guide-RNA line | This study |
| *UAS-mthl2-g* | UAS-guide-RNA line | This study |
| *UAS-mthl3-g* | UAS-guide-RNA line | This study |
| *UAS-mthl4-g* | UAS-guide-RNA line | This study |
| *UAS-mthl5-g* | UAS-guide-RNA line | This study |
| *UAS-mthl6-g* | UAS-guide-RNA line | This study |
| *UAS-mthl7-g* | UAS-guide-RNA line | This study |
| *UAS-mthl8-g* | UAS-guide-RNA line | This study |
| *UAS-mthl9-g* | UAS-guide-RNA line | This study |
| *UAS-mtt-g* | UAS-guide-RNA line | This study |
| *UAS-ninaE-g* | UAS-guide-RNA line | This study |
| *UAS-NPFR-g* | UAS-guide-RNA line | This study |
| *UAS-Oamb-g* | UAS-guide-RNA line | This study |
| *UAS-Octbeta1R-g* | UAS-guide-RNA line | This study |
| *UAS-Octbeta2R-g* | UAS-guide-RNA line | This study |
| *UAS-Octbeta3R-g* | UAS-guide-RNA line | This study |
| *UAS-Oct-TyrR-g* | UAS-guide-RNA line | This study |
| *UAS-PDFR-g* | UAS-guide-RNA line | This study |
| *UAS-PK1-R-g* | UAS-guide-RNA line | This study |
| *UAS-P2-R1-g* | UAS-guide-RNA line | This study |
| *UAS-P2-R2-g* | UAS-guide-RNA line | This study |
| *UAS-Proc-R-g* | UAS-guide-RNA line | This study |
| *UAS-rh2-g* | UAS-guide-RNA line | This study |
| *UAS-rh3-g* | UAS-guide-RNA line | This study |
| *UAS-rh4-g* | UAS-guide-RNA line | This study |
| *UAS-rh5-g* | UAS-guide-RNA line | This study |
| *UAS-rh6-g* | UAS-guide-RNA line | This study |

| | | |
|---|---|---|
| *UAS-rh7-g* | UAS-guide-RNA line | This study |
| *UAS-rho-g* | UAS-guide-RNA line | This study |
| *UAS-rho-4-g* | UAS-guide-RNA line | This study |
| *UAS-rho-5-g* | UAS-guide-RNA line | This study |
| *UAS-rho-6-g* | UAS-guide-RNA line | This study |
| *UAS-rho-7-g* | UAS-guide-RNA line | This study |
| *UAS-rk-g* | UAS-guide-RNA line | This study |
| *UAS-ru-g* | UAS-guide-RNA line | This study |
| *UAS-Rya-R-g* | UAS-guide-RNA line | This study |
| *UAS-smo-g* | UAS-guide-RNA line | This study |
| *UAS-smog-g* | UAS-guide-RNA line | This study |
| *UAS-sNPF-R-g* | UAS-guide-RNA line | This study |
| *UAS-SPR-g* | UAS-guide-RNA line | This study |
| *UAS-stan-g* | UAS-guide-RNA line | This study |
| *UAS-stet-g* | UAS-guide-RNA line | This study |
| *UAS-TkR86C-g* | UAS-guide-RNA line | This study |
| *UAS-TkR99D-g* | UAS-guide-RNA line | This study |
| *UAS-Tre1-g* | UAS-guide-RNA line | This study |
| *UAS-TrissinR-g* | UAS-guide-RNA line | This study |
| *UAS-TyrR* | UAS-guide-RNA line | This study |
| *UAS-TyRII* | UAS-guide-RNA line | This study |

Table S2: Primers used to generate UAS-guide lines.

| Gene name | # | Sequence |
|---|---|---|
| *CrzR* | 1 | GGCCCGGGTTCGATTCCCGGCCGATGCAAACATGATCGACGTGGGTGTGTTTCAGAGCTATGCTGGAAA |
| *CrzR* | 2 | GCGTGAAGGTGGCGTACATCTGCACCAGCCGGGAATCGAACC |
| *CrzR* | 3 | GATGTACGCCACCTTCACGCGTTTCAGAGCTATGCTGGAAAC |
| *CrzR* | 4 | TTTAACTTGCTATTTCTAGCTCTAAAACCGCCGCTCGACTTGCCCCCTTGCACCAGCCGGGAATCGAAC |
| *AkhR* | 1 | GGCCCGGGTTCGATTCCCGGCCGATGCAAAGGATATGGTCTTCAATGAGTTTCAGAGCTATGCTGGAAA |
| *AkhR* | 2 | TCACGTAGCTGGACAGATACTGCACCAGCCGGGAATCGAACC |
| *AkhR* | 3 | GTATCTGTCCAGCTACGTGAGTTTCAGAGCTATGCTGGAAAC |
| *AkhR* | 4 | TTTAACTTGCTATTTCTAGCTCTAAAACCGAATGTTGTAGAGTCCGTATGCACCAGCCGGGAATCGAAC |
| *CCAP-R* | 1 | GGCCCGGGTTCGATTCCCGGCCGATGCAAGTCACGACTTGGAGACGGGGTTTCAGAGCTATGCTGGAAA |
| *CCAP-R* | 2 | AATCGTCTTTACGATGATCGTGCACCAGCCGGGAATCGAACC |
| *CCAP-R* | 3 | CGATCATCGTAAAGACGATTGTTTCAGAGCTATGCTGGAAAC |
| *CCAP-R* | 4 | TTTAACTTGCTATTTCTAGCTCTAAAACACGACCAACCGTGTGGCAGCTGCACCAGCCGGGAATCGAAC |
| *CG30340* | 1 | GGCCCGGGTTCGATTCCCGGCCGATGCAGGGATGCGTGGGCTGCAAGCGTTTCAGAGCTATGCTGGAAA |
| *CG30340* | 2 | GGTGATCTCCCCATTGGGCTTGCACCAGCCGGGAATCGAACC |
| *CG30340* | 3 | AGCCCAATGGGGAGATCACCGTTTCAGAGCTATGCTGGAAAC |
| *CG30340* | 4 | TTTAACTTGCTATTTCTAGCTCTAAAACTTCTTCATAAAGGCACAGTATGCACCAGCCGGGAATCGAAC |
| *CG13995* | 1 | GGCCCGGGTTCGATTCCCGGCCGATGCAAGTGGTGGGAGAACTCGTATGTTTCAGAGCTATGCTGGAAA |
| *CG13995* | 2 | ATCGCGGATGCTTTACTGTGTGCACCAGCCGGGAATCGAACC |
| *CG13995* | 3 | CACAGTAAAGCATCCGCGATGTTTCAGAGCTATGCTGGAAAC |
| *CG13995* | 4 | TTTAACTTGCTATTTCTAGCTCTAAAACGCCTGCTCGGTTGATGCGCTTGCACCAGCCGGGAATCGAAC |
| *CG4313* | 1 | GGCCCGGGTTCGATTCCCGGCCGATGCATTTCCATAATGATACCCACCGTTTCAGAGCTATGCTGGAAA |
| *CG4313* | 2 | CTGGACGTGGTCACCTTCTCTGCACCAGCCGGGAATCGAACC |
| *CG4313* | 3 | GAGAAGGTGACCACGTCCAGGTTTCAGAGCTATGCTGGAAAC |
| *CG4313* | 4 | TTTAACTTGCTATTTCTAGCTCTAAAACTGTGAAGGAGGTGGTGCGCATGCACCAGCCGGGAATCGAAC |
| *moody* | 1 | GGCCCGGGTTCGATTCCCGGCCGATGCAGTTCCCCCGGCGGACGCGACGTTTCAGAGCTATGCTGGAAA |
| *moody* | 2 | CCACGTTGCGCACCTTGGGATGCACCAGCCGGGAATCGAACC |
| *moody* | 3 | TCCCAAGGTGCGCAACGTGGGTTTCAGAGCTATGCTGGAAAC |
| *moody* | 4 | TTTAACTTGCTATTTCTAGCTCTAAAACTTGGTCTTCCCGAAGGGCAGTGCACCAGCCGGGAATCGAAC |
| *Tre1* | 1 | GGCCCGGGTTCGATTCCCGGCCGATGCAGCCTGTGTCTTTGTGACGATGTTTCAGAGCTATGCTGGAAA |
| *Tre1* | 2 | ACCATGCTGAGGAGTGAAACTGCACCAGCCGGGAATCGAACC |

| | | |
|---|---|---|
| *Tre1* | 3 | GTTTCACTCCTCAGCATGGTGTTTCAGAGCTATGCTGGAAAC |
| *Tre1* | 4 | TTTAACTTGCTATTTCTAGCTCTAAAACTCATGACGGTCAGTCGATTGTGCACCAGCCGGGAATCGAAC |
| *CG15614* | 1 | GGCCCGGGTTCGATTCCCGGCCGATGCAGTTGTGCTCTTTGTGCTCTAGTTTCAGAGCTATGCTGGAAA |
| *CG15614* | 2 | CAACAATGATGCAGCGAGCCTGCACCAGCCGGGAATCGAACC |
| *CG15614* | 3 | GGCTCGCTGCATCATTGTTGGTTTCAGAGCTATGCTGGAAAC |
| *CG15614* | 4 | TTTAACTTGCTATTTCTAGCTCTAAAACATGACCAGAAATATGGCTGGGTGCACCAGCCGGGAATCGAAC |
| *CG7497* | 1 | GGCCCGGGTTCGATTCCCGGCCGATGCAGATTATTGGTGTTATTATCAGTTTCAGAGCTATGCTGGAAA |
| *CG7497* | 2 | GTAGAGCCACCAGATTGTTGTGCACCAGCCGGGAATCGAACC |
| *CG7497* | 3 | CAACAATCTGGTGGCTCTACGTTTCAGAGCTATGCTGGAAAC |
| *CG7497* | 4 | TTTAACTTGCTATTTCTAGCTCTAAAACATCACACAGAGCAAGGTGTGTGCACCAGCCGGGAATCGAAC |
| *FMRFaR* | 1 | GGCCCGGGTTCGATTCCCGGCCGATGCATGAGTGGTACAGCGGTTGCGGTTTCAGAGCTATGCTGGAAA |
| *FMRFaR* | 2 | ATAGGGAATACCGCCGGCGATGCACCAGCCGGGAATCGAACC |
| *FMRFaR* | 3 | TCGCCGGCGGTATTCCCTATGTTTCAGAGCTATGCTGGAAAC |
| *FMRFaR* | 4 | TTTAACTTGCTATTTCTAGCTCTAAAACGCCTGTATATCAGACAATTTTGCACCAGCCGGGAATCGAAC |
| *CNMaR* | 1 | GGCCCGGGTTCGATTCCCGGCCGATGCAGAGTATATAACTAGCAGTAGGTTTCAGAGCTATGCTGGAAA |
| *CNMaR* | 2 | GATGAGTTCGACGAGTTCGATGCACCAGCCGGGAATCGAACC |
| *CNMaR* | 3 | TCGAACTCGTCGAACTCATCGTTTCAGAGCTATGCTGGAAAC |
| *CNMaR* | 4 | TTTAACTTGCTATTTCTAGCTCTAAAACCCCGTTGAGCGGATGAAACCTGCACCAGCCGGGAATCGAAC |
| *Proc-R* | 1 | GGCCCGGGTTCGATTCCCGGCCGATGCAAGTATCATTTCAAGCTCTACGTTTCAGAGCTATGCTGGAAA |
| *Proc-R* | 2 | ATCGTAATTGTATGCTCGAATGCACCAGCCGGGAATCGAACC |
| *Proc-R* | 3 | TTCGAGCATACAATTACGATGTTTCAGAGCTATGCTGGAAAC |
| *Proc-R* | 4 | TTTAACTTGCTATTTCTAGCTCTAAAACGTGACGGCGATGTCGTGGAATGCACCAGCCGGGAATCGAAC |
| *CG33639* | 1 | GGCCCGGGTTCGATTCCCGGCCGATGCAACCTGAGGGACGACTTCTATGTTTCAGAGCTATGCTGGAAA |
| *CG33639* | 2 | GATAAAGCTCCAGGTGGGCGTGCACCAGCCGGGAATCGAACC |
| *CG33639* | 3 | CGCCCACCTGGAGCTTTATCGTTTCAGAGCTATGCTGGAAAC |
| *CG33639* | 4 | TTTAACTTGCTATTTCTAGCTCTAAAACGTAGACCATCATGATGCGCATGCACCAGCCGGGAATCGAAC |
| *SPR* | 1 | GGCCCGGGTTCGATTCCCGGCCGATGCACGTACTGTACCAGTACCGCTGTTTCAGAGCTATGCTGGAAA |
| *SPR* | 2 | AACGGAGATGGTGTGGCACATGCACCAGCCGGGAATCGAACC |
| *SPR* | 3 | TGTGCCACACCATCTCCGTTGTTTCAGAGCTATGCTGGAAAC |
| *SPR* | 4 | TTTAACTTGCTATTTCTAGCTCTAAAACCGTTCGGGCCATGGGGGCGTTGCACCAGCCGGGAATCGAAC |
| *CG13229* | 1 | GGCCCGGGTTCGATTCCCGGCCGATGCAAGTGCCAGGGCTATTGCCAGGTTTCAGAGCTATGCTGGAAA |
| *CG13229* | 2 | GCGTGACAGTCAATCCGATGTGCACCAGCCGGGAATCGAACC |

| | | |
|---|---|---|
| *CG13229* | 3 | CATCGGATTGACTGTCACGCGTTTCAGAGCTATGCTGGAAAC |
| *CG13229* | 4 | TTTAACTTGCTATTTCTAGCTCTAAAACCACGGCAGAAGTTTGATGAGTGCACCAGCCGGGAATCGAAC |
| *MsR2* | 1 | GGCCCGGGTTCGATTCCCGGCCGATGCAATGTCGCAGCCGCATTATTGGTTTCAGAGCTATGCTGGAAA |
| *MsR2* | 2 | AGGATACAGACAATCAGCGATGCACCAGCCGGGAATCGAACC |
| *MsR2* | 3 | TCGCTGATTGTCTGTATCCTGTTTCAGAGCTATGCTGGAAAC |
| *MsR2* | 4 | TTTAACTTGCTATTTCTAGCTCTAAAACACAGAGCACCGATGAGCCGCTGCACCAGCCGGGAATCGAAC |
| *MsR1* | 1 | GGCCCGGGTTCGATTCCCGGCCGATGCAGAAACTGAGCCGCTCTACTGGTTTCAGAGCTATGCTGGAAA |
| *MsR1* | 2 | GGTCGGCCACGGCCAGACCCTGCACCAGCCGGGAATCGAACC |
| *MsR1* | 3 | GGGTCTGGCCGTGGCCGACCGTTTCAGAGCTATGCTGGAAAC |
| *MsR1* | 4 | TTTAACTTGCTATTTCTAGCTCTAAAACGTAAGGCCAGGATCAATCGATGCACCAGCCGGGAATCGAAC |
| *AstC-R2* | 1 | GGCCCGGGTTCGATTCCCGGCCGATGCAACCAACGGCTGTGCCCATTCGTTTCAGAGCTATGCTGGAAA |
| *AstC-R2* | 2 | CCGACATGATCAACAGGAAGTGCACCAGCCGGGAATCGAACC |
| *AstC-R2* | 3 | CTTCCTGTTGATCATGTCGGGTTTCAGAGCTATGCTGGAAAC |
| *AstC-R2* | 4 | TTTAACTTGCTATTTCTAGCTCTAAAACTCGGAGCATTGTTGTTTCTCTGCACCAGCCGGGAATCGAAC |
| *AstC-R1* | 1 | GGCCCGGGTTCGATTCCCGGCCGATGCAGAACGAGAGCTTATATACCAGTTTCAGAGCTATGCTGGAAA |
| *AstC-R1* | 2 | GGCAATCGCTGAGACCACTTTGCACCAGCCGGGAATCGAACC |
| *AstC-R1* | 3 | AAGTGGTCTCAGCGATTGCCGTTTCAGAGCTATGCTGGAAAC |
| *AstC-R1* | 4 | TTTAACTTGCTATTTCTAGCTCTAAAACGCTTATTCATACAGGTAAAGTGCACCAGCCGGGAATCGAAC |
| *AstA-R2* | 1 | GGCCCGGGTTCGATTCCCGGCCGATGCAGGGATTCTTCGGCAACCTGCGTTTCAGAGCTATGCTGGAAA |
| *AstA-R2* | 2 | CATGATCATGCGCATGTAGATGCACCAGCCGGGAATCGAACC |
| *AstA-R2* | 3 | TCTACATGCGCATGATCATGGTTTCAGAGCTATGCTGGAAAC |
| *AstA-R2* | 4 | TTTAACTTGCTATTTCTAGCTCTAAAACGAAATTCTCGGAGAGGAAGGTGCACCAGCCGGGAATCGAAC |
| *AstA-R1* | 1 | GGCCCGGGTTCGATTCCCGGCCGATGCAAGCTGGCCAAAAGCCTCTTGGTTTCAGAGCTATGCTGGAAA |
| *AstA-R1* | 2 | AGACGGCCAGGTTGATTATCTGCACCAGCCGGGAATCGAACC |
| *AstA-R1* | 3 | GATAATCAACCTGGCCGTCTGTTTCAGAGCTATGCTGGAAAC |
| *AstA-R1* | 4 | TTTAACTTGCTATTTCTAGCTCTAAAACTGGTACTGATAAATCCTCACTGCACCAGCCGGGAATCGAAC |
| *CCHa2-R* | 1 | GGCCCGGGTTCGATTCCCGGCCGATGCATGTGCCCGTGCTGGACCGCCGTTTCAGAGCTATGCTGGAAA |
| *CCHa2-R* | 2 | GCTTTCCTGCGTGTAGACAATGCACCAGCCGGGAATCGAACC |
| *CCHa2-R* | 3 | TTGTCTACACGCAGGAAAGCGTTTCAGAGCTATGCTGGAAAC |
| *CCHa2-R* | 4 | TTTAACTTGCTATTTCTAGCTCTAAAACCTTGACGCAGATGCAGCAGATGCACCAGCCGGGAATCGAAC |
| *CCHa1-R* | 1 | GGCCCGGGTTCGATTCCCGGCCGATGCAGAGACACCCTACGTGCCCTAGTTTCAGAGCTATGCTGGAAA |
| *CCHa1-R* | 2 | ATCGCCGGATAATGCAGTCATGCACCAGCCGGGAATCGAACC |

| | | |
|---|---|---|
| *CCHa1-R* | 3 | TGACTGCATTATCCGGCGATGTTTCAGAGCTATGCTGGAAAC |
| *CCHa1-R* | 4 | TTTAACTTGCTATTTCTAGCTCTAAAACCTCACGAAGTATAGGGCCACTGCACCAGCCGGGAATCGAAC |
| *TrissinR* | 1 | GGCCCGGGTTCGATTCCCGGCCGATGCATAATGACGACGGCCACCACAGTTTCAGAGCTATGCTGGAAA |
| *TrissinR* | 2 | GGCAGGATGGCTGCTGCGGCTGCACCAGCCGGGAATCGAACC |
| *TrissinR* | 3 | GCCGCAGCAGCCATCCTGCCGTTTCAGAGCTATGCTGGAAAC |
| *TrissinR* | 4 | TTTAACTTGCTATTTCTAGCTCTAAAACGCCCGCTGGTGATGGGCGTGTGCACCAGCCGGGAATCGAAC |
| *PK2-R2* | 1 | GGCCCGGGTTCGATTCCCGGCCGATGCACAACCTGACCAGCCTTCTCCGTTTCAGAGCTATGCTGGAAA |
| *PK2-R2* | 2 | TTTCCGAGAGAACGCTCTCCTGCACCAGCCGGGAATCGAACC |
| *PK2-R2* | 3 | GGAGAGCGTTCTCTCGGAAAGTTTCAGAGCTATGCTGGAAAC |
| *PK2-R2* | 4 | TTTAACTTGCTATTTCTAGCTCTAAAACGATGACTCGCGTTTGGGCGCTGCACCAGCCGGGAATCGAAC |
| *PK2-R1* | 1 | GGCCCGGGTTCGATTCCCGGCCGATGCACCCAATCAGAGTCCCTCCATGTTTCAGAGCTATGCTGGAAA |
| *PK2-R1* | 2 | TCGTTCTGGTAGACCACCGATGCACCAGCCGGGAATCGAACC |
| *PK2-R1* | 3 | TCGGTGGTCTACCAGAACGAGTTTCAGAGCTATGCTGGAAAC |
| *PK2-R1* | 4 | TTTAACTTGCTATTTCTAGCTCTAAAACGACCAAACTGTCGCGTCAGATGCACCAGCCGGGAATCGAAC |
| *PK1-R* | 1 | GGCCCGGGTTCGATTCCCGGCCGATGCACCAAGTACCCGTACGTGTTTGTTTCAGAGCTATGCTGGAAA |
| *PK1-R* | 2 | CCCGTGCAGGGGCGTAGATGTGCACCAGCCGGGAATCGAACC |
| *PK1-R* | 3 | CATCTACGCCCCTGCACGGGGTTTCAGAGCTATGCTGGAAAC |
| *PK1-R* | 4 | TTTAACTTGCTATTTCTAGCTCTAAAACTCCTCAGGCGGCGCGATTCGTGCACCAGCCGGGAATCGAAC |
| *CapaR* | 1 | GGCCCGGGTTCGATTCCCGGCCGATGCAGGAGAGGAGGACTATGCCTGGTTTCAGAGCTATGCTGGAAA |
| *CapaR* | 2 | GTTATAAATATGCCACCAAATGCACCAGCCGGGAATCGAACC |
| *CapaR* | 3 | TTTGGTGGCATATTTATAACGTTTCAGAGCTATGCTGGAAAC |
| *CapaR* | 4 | TTTAACTTGCTATTTCTAGCTCTAAAACAAGAGATCCGGATACTGATGTGCACCAGCCGGGAATCGAAC |
| *ETHR* | 1 | GGCCCGGGTTCGATTCCCGGCCGATGCAATCATGCTGCTGGGTGTGGTGTTTCAGAGCTATGCTGGAAA |
| *ETHR* | 2 | TCGTCTTCACAATCACGATGTGCACCAGCCGGGAATCGAACC |
| *ETHR* | 3 | CATCGTGATTGTGAAGACGAGTTTCAGAGCTATGCTGGAAAC |
| *ETHR* | 4 | TTTAACTTGCTATTTCTAGCTCTAAAACTGGTGCAGACGTAGCCGGCCTGCACCAGCCGGGAATCGAAC |
| *TkR99D* | 1 | GGCCCGGGTTCGATTCCCGGCCGATGCAAGCAACTGGAGCACCCCCGCGTTTCAGAGCTATGCTGGAAA |
| *TkR99D* | 2 | CCCGTCGCCACAATGACCATTGCACCAGCCGGGAATCGAACC |
| *TkR99D* | 3 | ATGGTCATTGTGGCGACGGGGTTTCAGAGCTATGCTGGAAAC |
| *TkR99D* | 4 | TTTAACTTGCTATTTCTAGCTCTAAAACAAGGGCCAGTCGCTATCCAGTGCACCAGCCGGGAATCGAAC |
| *TkR86C* | 1 | GGCCCGGGTTCGATTCCCGGCCGATGCAGTCGAGCATTATCGACAATCGTTTCAGAGCTATGCTGGAAA |
| *TkR86C* | 2 | ATCTGAGTTCAGCATGAATATGCACCAGCCGGGAATCGAACC |

| | | |
|---|---|---|
| *TkR86C* | 3 | TATTCATGCTGAACTCAGATGTTTCAGAGCTATGCTGGAAAC |
| *TkR86C* | 4 | TTTAACTTGCTATTTCTAGCTCTAAAACATTGATCTACTGCCCCACAGTGCACCAGCCGGGAATCGAAC |
| *Lkr* | 1 | GGCCCGGGTTCGATTCCCGGCCGATGCAGGAATTCCTGCCCGGAGCCGGTTTCAGAGCTATGCTGGAAA |
| *Lkr* | 2 | GGTCGTGGCCACCACCCAGATGCACCAGCCGGGAATCGAACC |
| *Lkr* | 3 | TCTGGGTGGTGGCCACGACCGTTTCAGAGCTATGCTGGAAAC |
| *Lkr* | 4 | TTTAACTTGCTATTTCTAGCTCTAAAACGGACGAAGGGGCAGAAGCTGTGCACCAGCCGGGAATCGAAC |
| *Rya-R* | 1 | GGCCCGGGTTCGATTCCCGGCCGATGCAGCAGCCGATGCTGCGGAACGGTTTCAGAGCTATGCTGGAAA |
| *Rya-R* | 2 | GCGAGTAGTTCACAAAGTGATGCACCAGCCGGGAATCGAACC |
| *Rya-R* | 3 | TCACTTTGTGAACTACTCGCGTTTCAGAGCTATGCTGGAAAC |
| *Rya-R* | 4 | TTTAACTTGCTATTTCTAGCTCTAAAACAGCGGCACGACGAACTGCAGTGCACCAGCCGGGAATCGAAC |
| *SIFaR* | 1 | GGCCCGGGTTCGATTCCCGGCCGATGCACCATTCCGACCATGGCGCCGGTTTCAGAGCTATGCTGGAAA |
| *SIFaR* | 2 | AGGCTGCCACGGAGACACCTTGCACCAGCCGGGAATCGAACC |
| *SIFaR* | 3 | AGGTGTCTCCGTGGCAGCCTGTTTCAGAGCTATGCTGGAAAC |
| *SIFaR* | 4 | TTTAACTTGCTATTTCTAGCTCTAAAACGGGCGTCGGAGAAGACCTCCTGCACCAGCCGGGAATCGAAC |
| *CCKLR-17D1* | 1 | GGCCCGGGTTCGATTCCCGGCCGATGCACGCGGAACCCGAACTCCATAGTTTCAGAGCTATGCTGGAAA |
| *CCKLR-17D1* | 2 | GGTGCGCGACCTCAGCGGGTTGCACCAGCCGGGAATCGAACC |
| *CCKLR-17D1* | 3 | ACCCGCTGAGGTCGCGCACCGTTTCAGAGCTATGCTGGAAAC |
| *CCKLR-17D1* | 4 | TTTAACTTGCTATTTCTAGCTCTAAAACCCGGCCCACTGCTGCCAAAGTGCACCAGCCGGGAATCGAAC |
| *CCKLR-17D3* | 1 | GGCCCGGGTTCGATTCCCGGCCGATGCATATGGCGATGATGATAGGGAGTTTCAGAGCTATGCTGGAAA |
| *CCKLR-17D3* | 2 | AGGTCCAGGACGAAACGGCCTGCACCAGCCGGGAATCGAACC |
| *CCKLR-17D3* | 3 | GGCCGTTTCGTCCTGGACCTGTTTCAGAGCTATGCTGGAAAC |
| *CCKLR-17D3* | 4 | TTTAACTTGCTATTTCTAGCTCTAAAACCGCAGAGGACGAGAAGCGGCTGCACCAGCCGGGAATCGAAC |
| *sNPF-R* | 1 | GGCCCGGGTTCGATTCCCGGCCGATGCAGGGAACTACTAGCGCTATCTGTTTCAGAGCTATGCTGGAAA |
| *sNPF-R* | 2 | TGAACGTGTAAAGCGGAGTATGCACCAGCCGGGAATCGAACC |
| *sNPF-R* | 3 | TACTCCGCTTTACACGTTCAGTTTCAGAGCTATGCTGGAAAC |
| *sNPF-R* | 4 | TTTAACTTGCTATTTCTAGCTCTAAAACCCTTGCCATGATGCAGTCGCTGCACCAGCCGGGAATCGAAC |
| *NPFR* | 1 | GGCCCGGGTTCGATTCCCGGCCGATGCAGCAGATGGGGAGCATCTGAGGTTTCAGAGCTATGCTGGAAA |
| *NPFR* | 2 | GCGGCCTGCTCCCAGTCTCATGCACCAGCCGGGAATCGAACC |
| *NPFR* | 3 | TGAGACTGGGAGCAGGCCGCGTTTCAGAGCTATGCTGGAAAC |
| *NPFR* | 4 | TTTAACTTGCTATTTCTAGCTCTAAAACTAGGGCCAGTACTTGGACAGTGCACCAGCCGGGAATCGAAC |
| *Dh31-R* | 1 | GGCCCGGGTTCGATTCCCGGCCGATGCATTCCGCCTACGAACTCTGCCGTTTCAGAGCTATGCTGGAAA |
| *Dh31-R* | 2 | GCCTCTTCTCGGAAATGAATTGCACCAGCCGGGAATCGAACC |

| | | |
|---|---|---|
| *Dh31-R* | 3 | ATTCATTTCCGAGAAGAGGCGTTTCAGAGCTATGCTGGAAAC |
| *Dh31-R* | 4 | TTTAACTTGCTATTTCTAGCTCTAAAACTGAATACTGGCCGGGGCATTTGCACCAGCCGGGAATCGAAC |
| *hec* | 1 | GGCCCGGGTTCGATTCCCGGCCGATGCAAAATGTGGATGTGGCATCGCGTTTCAGAGCTATGCTGGAAA |
| *hec* | 2 | CCAGGTCCTCATAGTCCACGTGCACCAGCCGGGAATCGAACC |
| *hec* | 3 | CGTGGACTATGAGGACCTGGGTTTCAGAGCTATGCTGGAAAC |
| *hec* | 4 | TTTAACTTGCTATTTCTAGCTCTAAAACGCAGGGAGAGGGCATAACCCTGCACCAGCCGGGAATCGAAC |
| *Dh44-R2* | 1 | GGCCCGGGTTCGATTCCCGGCCGATGCAAGACGACGATTTGAGGGCACGTTTCAGAGCTATGCTGGAAA |
| *Dh44-R2* | 2 | GACTGCCGGCGTTTGTGCGTTGCACCAGCCGGGAATCGAACC |
| *Dh44-R2* | 3 | ACGCACAAACGCCGGCAGTCGTTTCAGAGCTATGCTGGAAAC |
| *Dh44-R2* | 4 | TTTAACTTGCTATTTCTAGCTCTAAAACACGTTCCGTTTGGAAAGCAATGCACCAGCCGGGAATCGAAC |
| *Dh44-R1* | 1 | GGCCCGGGTTCGATTCCCGGCCGATGCATCGATTCGGTGAACGCCAGCGTTTCAGAGCTATGCTGGAAA |
| *Dh44-R1* | 2 | TTGGCGTGGCAGAACCTCGTTGCACCAGCCGGGAATCGAACC |
| *Dh44-R1* | 3 | ACGAGGTTCTGCCACGCCAAGTTTCAGAGCTATGCTGGAAAC |
| *Dh44-R1* | 4 | TTTAACTTGCTATTTCTAGCTCTAAAACCTGTATTGGCCGAGCGCAGCTGCACCAGCCGGGAATCGAAC |
| *PDFR* | 1 | GGCCCGGGTTCGATTCCCGGCCGATGCATCGAACATTCTCGACTGCGGGTTTCAGAGCTATGCTGGAAA |
| *PDFR* | 2 | GCCGGAGTGGGTGGCCAGCATGCACCAGCCGGGAATCGAACC |
| *PDFR* | 3 | TGCTGGCCACCCACTCCGGCGTTTCAGAGCTATGCTGGAAAC |
| *PDFR* | 4 | TTTAACTTGCTATTTCTAGCTCTAAAACCCTGGCAATGTCTATGTAGGTGCACCAGCCGGGAATCGAAC |
| *CG11318* | 1 | GGCCCGGGTTCGATTCCCGGCCGATGCATGGAGAGGAACGTAATTCATGTTTCAGAGCTATGCTGGAAA |
| *CG11318* | 2 | TGTCCTGCTGCATCAGGGCGTGCACCAGCCGGGAATCGAACC |
| *CG11318* | 3 | CGCCCTGATGCAGCAGGACAGTTTCAGAGCTATGCTGGAAAC |
| *CG11318* | 4 | TTTAACTTGCTATTTCTAGCTCTAAAACTCAGTTGACCAGGTTTCATATGCACCAGCCGGGAATCGAAC |
| *CG15556* | 1 | GGCCCGGGTTCGATTCCCGGCCGATGCAGCTGTCGCACAATCCCTACAGTTTCAGAGCTATGCTGGAAA |
| *CG15556* | 2 | GAGTTTGAGTCAGGAGGTTATGCACCAGCCGGGAATCGAACC |
| *CG15556* | 3 | TAACCTCCTGACTCAAACTCGTTTCAGAGCTATGCTGGAAAC |
| *CG15556* | 4 | TTTAACTTGCTATTTCTAGCTCTAAAACTCGTTTGATTTCCTCTTCCGTGCACCAGCCGGGAATCGAAC |
| *5-HT1B* | 1 | GGCCCGGGTTCGATTCCCGGCCGATGCAATCGCCATGGCCGTGGTGCTGTTTCAGAGCTATGCTGGAAA |
| *5-HT1B* | 2 | GGCAATAATGTAGATTTTCCTGCACCAGCCGGGAATCGAACC |
| *5-HT1B* | 3 | GGAAAATCTACATTATTGCCGTTTCAGAGCTATGCTGGAAAC |
| *5-HT1B* | 4 | TTTAACTTGCTATTTCTAGCTCTAAAACCCCGTCCCGTCGCCGTCCACTGCACCAGCCGGGAATCGAAC |
| *5-HT1A* | 1 | GGCCCGGGTTCGATTCCCGGCCGATGCATAGCGAACAGCATGAATGACGTTTCAGAGCTATGCTGGAAA |
| *5-HT1A* | 2 | GGATAATGGCCGCTATGACATGCACCAGCCGGGAATCGAACC |

| | | |
|---|---|---|
| *5-HT1A* | 3 | TGTCATAGCGGCCATTATCCGTTTCAGAGCTATGCTGGAAAC |
| *5-HT1A* | 4 | TTTAACTTGCTATTTCTAGCTCTAAAACCATCGACGGGTCGCGGTCGTTGCACCAGCCGGGAATCGAAC |
| *5-HT7* | 1 | GGCCCGGGTTCGATTCCCGGCCGATGCACACAGAAACCACAGAACCCAGTTTCAGAGCTATGCTGGAAA |
| *5-HT7* | 2 | AAATTGCTGCTGGTGATGCTGCACCAGCCGGGAATCGAACC |
| *5-HT7* | 3 | GCATCACCAGCAGCAATTTGTTTCAGAGCTATGCTGGAAAC |
| *5-HT7* | 4 | TTTAACTTGCTATTTCTAGCTCTAAAACGAAGAGCCACACAGAGATCCTGCACCAGCCGGGAATCGAAC |
| *mAChR-A* | 1 | GGCCCGGGTTCGATTCCCGGCCGATGCAAAAGGGACCTCGGTACTCACGTTTCAGAGCTATGCTGGAAA |
| *mAChR-A* | 2 | CGTCCGGTGAACTCTCGGAGTGCACCAGCCGGGAATCGAACC |
| *mAChR-A* | 3 | CTCCGAGAGTTCACCGGACGGTTTCAGAGCTATGCTGGAAAC |
| *mAChR-A* | 4 | TTTAACTTGCTATTTCTAGCTCTAAAACGCTCGTAGGCGAAGTCGTCGTGCACCAGCCGGGAATCGAAC |
| *mAChR-B* | 1 | GGCCCGGGTTCGATTCCCGGCCGATGCAGCGGTCGCTTAACAAGTCGGGTTTCAGAGCTATGCTGGAAA |
| *mAChR-B* | 2 | TGTGCGTCAGGACTCGCTTGTGCACCAGCCGGGAATCGAACC |
| *mAChR-B* | 3 | CAAGCGAGTCCTGACGCACAGTTTCAGAGCTATGCTGGAAAC |
| *mAChR-B* | 4 | TTTAACTTGCTATTTCTAGCTCTAAAACTGAGCACGTATATCGTGTAGTGCACCAGCCGGGAATCGAAC |
| *Oct-TyrR* | 1 | GGCCCGGGTTCGATTCCCGGCCGATGCAGTTTGTAAATGTCACCACAAGTTTCAGAGCTATGCTGGAAA |
| *Oct-TyrR* | 2 | AGAGAACCAGGGCGGTGAGATGCACCAGCCGGGAATCGAACC |
| *Oct-TyrR* | 3 | TCTCACCGCCCTGGTTCTCTGTTTCAGAGCTATGCTGGAAAC |
| *Oct-TyrR* | 4 | TTTAACTTGCTATTTCTAGCTCTAAAACGCACGACCACGTGCTGCTGTTGCACCAGCCGGGAATCGAAC |
| *Oamb* | 1 | GGCCCGGGTTCGATTCCCGGCCGATGCAACCAGCCAATCTGATCTCCCGTTTCAGAGCTATGCTGGAAA |
| *Oamb* | 2 | CCACATAGCGGTCCAGTGATTGCACCAGCCGGGAATCGAACC |
| *Oamb* | 3 | ATCACTGGACCGCTATGTGGGTTTCAGAGCTATGCTGGAAAC |
| *Oamb* | 4 | TTTAACTTGCTATTTCTAGCTCTAAAACCGCCCTGGTCCTTAATGCTATGCACCAGCCGGGAATCGAAC |
| *Dop1R2* | 1 | GGCCCGGGTTCGATTCCCGGCCGATGCACCTCCTGGCGCTCCTCCGGGGTTTCAGAGCTATGCTGGAAA |
| *Dop1R2* | 2 | GATAGCCTAGGTGCTCAGTGTGCACCAGCCGGGAATCGAACC |
| *Dop1R2* | 3 | CACTGAGCACCTAGGCTATCGTTTCAGAGCTATGCTGGAAAC |
| *Dop1R2* | 4 | TTTAACTTGCTATTTCTAGCTCTAAAACGCCGAGCCAGGTGACGATTGTGCACCAGCCGGGAATCGAAC |
| *CG18208* | 1 | GGCCCGGGTTCGATTCCCGGCCGATGCACATCAGCACCGATGACTTTCGTTTCAGAGCTATGCTGGAAA |
| *CG18208* | 2 | GTAGCCCATTAGCTCATTGGTGCACCAGCCGGGAATCGAACC |
| *CG18208* | 3 | CCAATGAGCTAATGGGCTACGTTTCAGAGCTATGCTGGAAAC |
| *CG18208* | 4 | TTTAACTTGCTATTTCTAGCTCTAAAACCGCCCGCGCCTTGGCGGCAATGCACCAGCCGGGAATCGAAC |
| *TyrRII* | 1 | GGCCCGGGTTCGATTCCCGGCCGATGCATGTTGTCAACAGCCCTCACTGGTTTCAGAGCTATGCTGGAAA |
| *TyrRII* | 2 | CTGCAGAGCAGGATGTCCAGTGCACCAGCCGGGAATCGAACC |

| | | |
|---|---|---|
| *TyrRII* | 3 | CTGGACATCCTGCTCTGCAGGTTTCAGAGCTATGCTGGAAAC |
| *TyrRII* | 4 | TTTAACTTGCTATTTCTAGCTCTAAAACCACGACGAGAATCATGAGGATGCACCAGCCGGGAATCGAAC |
| *TyrR* | 1 | GGCCCGGGTTCGATTCCCGGCCGATGCATTATCCGTCGCTCATGTATGGTTTCAGAGCTATGCTGGAAA |
| *TyrR* | 2 | CGAGAAGGTCGGCCACCGCCTGCACCAGCCGGGAATCGAACC |
| *TyrR* | 3 | GGCGGTGGCCGACCTTCTCGGTTTCAGAGCTATGCTGGAAAC |
| *TyrR* | 4 | TTTAACTTGCTATTTCTAGCTCTAAAACATCGTTTGGAGCGCCGTTTATGCACCAGCCGGGAATCGAAC |
| *Dop2R* | 1 | GGCCCGGGTTCGATTCCCGGCCGATGCAGGCGGCAACGGGAGCAACTGGTTTCAGAGCTATGCTGGAAA |
| *Dop2R* | 2 | AAACCGTGCTCCCATTGTAGTGCACCAGCCGGGAATCGAACC |
| *Dop2R* | 3 | CTACAATGGGAGCACGGTTTGTTTCAGAGCTATGCTGGAAAC |
| *Dop2R* | 4 | TTTAACTTGCTATTTCTAGCTCTAAAACAAGAGGGTCAGGATGGGGAATGCACCAGCCGGGAATCGAAC |
| *Dop1R1* | 1 | GGCCCGGGTTCGATTCCCGGCCGATGCAAATCACACCAGTGGGAGCACGTTTCAGAGCTATGCTGGAAA |
| *Dop1R1* | 2 | ACTATTGTATCCATTTCAGATGCACCAGCCGGGAATCGAACC |
| *Dop1R1* | 3 | TCTGAAATGGATACAATAGTGTTTCAGAGCTATGCTGGAAAC |
| *Dop1R1* | 4 | TTTAACTTGCTATTTCTAGCTCTAAAACGCGCAGGCTGCGCTCCGTGTTGCACCAGCCGGGAATCGAAC |
| *Octbeta2R* | 1 | GGCCCGGGTTCGATTCCCGGCCGATGCAGCCCGGAGCCACCGCGGCAAGTTTCAGAGCTATGCTGGAAA |
| *Octbeta2R* | 2 | GTTACTTGCACACTAAAGTTTGCACCAGCCGGGAATCGAACC |
| *Octbeta2R* | 3 | AACTTTAGTGTGCAAGTAACGTTTCAGAGCTATGCTGGAAAC |
| *Octbeta2R* | 4 | TTTAACTTGCTATTTCTAGCTCTAAAACCAAATCGCACAGGAAGGGGCTGCACCAGCCGGGAATCGAAC |
| *Octbeta1R* | 1 | GGCCCGGGTTCGATTCCCGGCCGATGCACACGACCAGGACAATACTGGGTTTCAGAGCTATGCTGGAAA |
| *Octbeta1R* | 2 | AGATCATGACGGAAGCATTATGCACCAGCCGGGAATCGAACC |
| *Octbeta1R* | 3 | TAATGCTTCCGTCATGATCTGTTTCAGAGCTATGCTGGAAAC |
| *Octbeta1R* | 4 | TTTAACTTGCTATTTCTAGCTCTAAAACCACCATCAATAGCATGATGATGCACCAGCCGGGAATCGAAC |
| *Octbeta3R* | 1 | GGCCCGGGTTCGATTCCCGGCCGATGCAGCCTATAACATCGCAATTTGGTTTCAGAGCTATGCTGGAAA |
| *Octbeta3R* | 2 | GGCTGTTGTACACGTTGCACTGCACCAGCCGGGAATCGAACC |
| *Octbeta3R* | 3 | GTGCAACGTGTACAACAGCCGTTTCAGAGCTATGCTGGAAAC |
| *Octbeta3R* | 4 | TTTAACTTGCTATTTCTAGCTCTAAAACGATGCAGGGAAATCTCCCGCTGCACCAGCCGGGAATCGAAC |
| *5-HT2B* | 1 | GGCCCGGGTTCGATTCCCGGCCGATGCAGAGGATGTGTATGCCTCGCTGTTTCAGAGCTATGCTGGAAA |
| *5-HT2B* | 2 | CGGAGATGGTGCACAGGTGCTGCACCAGCCGGGAATCGAACC |
| *5-HT2B* | 3 | GCACCTGTGCACCATCTCCGGTTTCAGAGCTATGCTGGAAAC |
| *5-HT2B* | 4 | TTTAACTTGCTATTTCTAGCTCTAAAACGTATCTGGCAAGTTCCATTCTGCACCAGCCGGGAATCGAAC |
| *5-HT2A* | 1 | GGCCCGGGTTCGATTCCCGGCCGATGCACAGCCGTTTATTGACTGCGAGTTTCAGAGCTATGCTGGAAA |
| *5-HT2A* | 2 | TGGGGACTGTAGTCCAGAATGCACCAGCCGGGAATCGAACC |

| | | |
|---|---|---|
| *5-HT2A* | 3 | TTCTGGACTACAGTCCCCAGTTTCAGAGCTATGCTGGAAAC |
| *5-HT2A* | 4 | TTTAACTTGCTATTTCTAGCTCTAAAACAACTGATGAAGCACATGTGCTGCACCAGCCGGGAATCGAAC |
| *AdoR* | 1 | GGCCCGGGTTCGATTCCCGGCCGATGCAGATCACCGATTTCTCCTTCGGTTTCAGAGCTATGCTGGAAA |
| *AdoR* | 2 | GGGATACTATGTAGTAGTTGTGCACCAGCCGGGAATCGAACC |
| *AdoR* | 3 | CAACTACTACATAGTATCCCGTTTCAGAGCTATGCTGGAAAC |
| *AdoR* | 4 | TTTAACTTGCTATTTCTAGCTCTAAAACTGCGGGTGCGGACATTCCTTTGCACCAGCCGGGAATCGAAC |
| *mAchR-C* | 1 | GGCCCGGGTTCGATTCCCGGCCGATGCATGGGTGTCGTCCCGTTCCTGGTTTCAGAGCTATGCTGGAAA |
| *mAchR-C* | 2 | TCTATAGTGCAGAGCATAGATGCACCAGCCGGGAATCGAACC |
| *mAchR-C* | 3 | TCTATGCTCTGCACTATAGAGTTTCAGAGCTATGCTGGAAAC |
| *mAchR-C* | 4 | TTTAACTTGCTATTTCTAGCTCTAAAACGTCTGGATGGAGCAGCAGATTGCACCAGCCGGGAATCGAAC |
| *CG13579* | 1 | GGCCCGGGTTCGATTCCCGGCCGATGCATGCTCGCATCCGCGCTACTCGTTTCAGAGCTATGCTGGAAA |
| *CG13579* | 2 | TGCAAACCAGGATGACGGCGTGCACCAGCCGGGAATCGAACC |
| *CG13579* | 3 | CGCCGTCATCCTGGTTTGCAGTTTCAGAGCTATGCTGGAAAC |
| *CG13579* | 4 | TTTAACTTGCTATTTCTAGCTCTAAAACGAAGCGGCTCATGTGGAAGCTGCACCAGCCGGGAATCGAAC |
| *DopEcR* | 1 | GGCCCGGGTTCGATTCCCGGCCGATGCAGCTCATATCGATACTGGGCGGTTTCAGAGCTATGCTGGAAA |
| *DopEcR* | 2 | GTCAAAGCGGGATACACGGATGCACCAGCCGGGAATCGAACC |
| *DopEcR* | 3 | TCCGTGTATCCCGCTTTGACGTTTCAGAGCTATGCTGGAAAC |
| *DopEcR* | 4 | TTTAACTTGCTATTTCTAGCTCTAAAACTATTCCCCCAGTCCAGCATGTGCACCAGCCGGGAATCGAAC |
| *Cirl* | 1 | GGCCCGGGTTCGATTCCCGGCCGATGCACAAGCAGAGCTGCGGCGTGTGTTTCAGAGCTATGCTGGAAA |
| *Cirl* | 2 | AATCTGGGAAGACCACGCTGTGCACCAGCCGGGAATCGAACC |
| *Cirl* | 3 | CAGCGTGGTCTTCCCAGATTGTTTCAGAGCTATGCTGGAAAC |
| *Cirl* | 4 | TTTAACTTGCTATTTCTAGCTCTAAAACCGCGTTGACATCGCTGTCGTTGCACCAGCCGGGAATCGAAC |
| *stan* | 1 | GGCCCGGGTTCGATTCCCGGCCGATGCAAAACGAGGGAGTTCCCTCAAGTTTCAGAGCTATGCTGGAAA |
| *stan* | 2 | TGCTCTGGTGCTCGATGGCATGCACCAGCCGGGAATCGAACC |
| *stan* | 3 | TGCCATCGAGCACCAGAGCAGTTTCAGAGCTATGCTGGAAAC |
| *stan* | 4 | TTTAACTTGCTATTTCTAGCTCTAAAACACGGGAACATTCTCTAGCACTGCACCAGCCGGGAATCGAAC |
| *mthl14* | 1 | GGCCCGGGTTCGATTCCCGGCCGATGCAACTGTAAATAACAGTTCGAAGTTTCAGAGCTATGCTGGAAA |
| *mthl14* | 2 | TCACAGTGTATGATTCGTTGTGCACCAGCCGGGAATCGAACC |
| *mthl14* | 3 | CAACGAATCATACACTGTGAGTTTCAGAGCTATGCTGGAAAC |
| *mthl14* | 4 | TTTAACTTGCTATTTCTAGCTCTAAAACAGCACAGAGACATCAAACTTTGCACCAGCCGGGAATCGAAC |
| *mthl5* | 1 | GGCCCGGGTTCGATTCCCGGCCGATGCAGTCATCGGGATTCCAAATTGGTTTCAGAGCTATGCTGGAAA |
| *mthl5* | 2 | CATAATCCGATTGTATTCCATGCACCAGCCGGGAATCGAACC |

| | | |
|---|---|---|
| *mthl5* | 3 | TGGAATACAATCGGATTATGGTTTCAGAGCTATGCTGGAAAC |
| *mthl5* | 4 | TTTAACTTGCTATTTCTAGCTCTAAAACGCCGTCCGTGACTCGCAGGATGCACCAGCCGGGAATCGAAC |
| *mthl9* | 1 | GGCCCGGGTTCGATTCCCGGCCGATGCATCCCTGTGCCTACGCCCACAGTTTCAGAGCTATGCTGGAAA |
| *mthl9* | 2 | GCAACCCAGCTCTGTGCGCATGCACCAGCCGGGAATCGAACC |
| *mthl9* | 3 | TGCGCACAGAGCTGGGTTGCGTTTCAGAGCTATGCTGGAAAC |
| *mthl9* | 4 | TTTAACTTGCTATTTCTAGCTCTAAAACACGATCATGGACAGCAAGTATGCACCAGCCGGGAATCGAAC |
| *mthl8* | 1 | GGCCCGGGTTCGATTCCCGGCCGATGCATTCTGGAACACACTGCTCGTGTTTCAGAGCTATGCTGGAAA |
| *mthl8* | 2 | ATGGAGCAGCGGAGTGAAGCTGCACCAGCCGGGAATCGAACC |
| *mthl8* | 3 | GCTTCACTCCGCTGCTCCATGTTTCAGAGCTATGCTGGAAAC |
| *mthl8* | 4 | TTTAACTTGCTATTTCTAGCTCTAAAACTCAGCACGATAAACATTGATTGCACCAGCCGGGAATCGAAC |
| *mthl1* | 1 | GGCCCGGGTTCGATTCCCGGCCGATGCAGTCGATGAGTTTGGCCATCGGTTTCAGAGCTATGCTGGAAA |
| *mthl1* | 2 | CACATAGAGCGTGCCATTGGTGCACCAGCCGGGAATCGAACC |
| *mthl1* | 3 | CCAATGGCACGCTCTATGTGGTTTCAGAGCTATGCTGGAAAC |
| *mthl1* | 4 | TTTAACTTGCTATTTCTAGCTCTAAAACAAGTGGCGGACAGGAAGACATGCACCAGCCGGGAATCGAAC |
| *mthl15* | 1 | GGCCCGGGTTCGATTCCCGGCCGATGCATCGCCGCACTGTGGACAATGGTTTCAGAGCTATGCTGGAAA |
| *mthl15* | 2 | GGTTTGCATGTATTATTTTCTGCACCAGCCGGGAATCGAACC |
| *mthl15* | 3 | GAAAATAATACATGCAAACCGTTTCAGAGCTATGCTGGAAAC |
| *mthl15* | 4 | TTTAACTTGCTATTTCTAGCTCTAAAACTCGGCACAACAACGACGAATTGCACCAGCCGGGAATCGAAC |
| *mthl13* | 1 | GGCCCGGGTTCGATTCCCGGCCGATGCACGGAAAAATATGATGGCCAAGTTTCAGAGCTATGCTGGAAA |
| *mthl13* | 2 | TAAGCCAGATTAGATAGTCATGCACCAGCCGGGAATCGAACC |
| *mthl13* | 3 | TGACTATCTAATCTGGCTTAGTTTCAGAGCTATGCTGGAAAC |
| *mthl13* | 4 | TTTAACTTGCTATTTCTAGCTCTAAAACTATGCAATAGCTGATTATGGTGCACCAGCCGGGAATCGAAC |
| *mthl10* | 1 | GGCCCGGGTTCGATTCCCGGCCGATGCATCCCTTCGGCACTTACGTTAGTTTCAGAGCTATGCTGGAAA |
| *mthl10* | 2 | ATATCGTAGACTCGTTGCGGTGCACCAGCCGGGAATCGAACC |
| *mthl10* | 3 | CCGCAACGAGTCTACGATATGTTTCAGAGCTATGCTGGAAAC |
| *mthl10* | 4 | TTTAACTTGCTATTTCTAGCTCTAAAACACGGAGAGCCCTATTAAATATGCACCAGCCGGGAATCGAAC |
| *mthl11* | 1 | GGCCCGGGTTCGATTCCCGGCCGATGCACTCTTAGGTATCCTTGTGATGTTTCAGAGCTATGCTGGAAA |
| *mthl11* | 2 | GGGTGTATTCGTAGGTAAGGTGCACCAGCCGGGAATCGAACC |
| *mthl11* | 3 | CCTTACCTACGAATACACCCGTTTCAGAGCTATGCTGGAAAC |
| *mthl11* | 4 | TTTAACTTGCTATTTCTAGCTCTAAAACTTACAGTGAAGTAGCCAGCATGCACCAGCCGGGAATCGAAC |
| *mthl6* | 1 | GGCCCGGGTTCGATTCCCGGCCGATGCACAATCGGAGGCAGTTATTCCGTTTCAGAGCTATGCTGGAAA |
| *mthl6* | 2 | GTGTGATCTTCAAGTAGGGATGCACCAGCCGGGAATCGAACC |

| | | |
|---|---|---|
| *mthl6* | 3 | TCCCTACTTGAAGATCACACGTTTCAGAGCTATGCTGGAAAC |
| *mthl6* | 4 | TTTAACTTGCTATTTCTAGCTCTAAAACTATTTGATGACTCATAACAGTGCACCAGCCGGGAATCGAAC |
| *mthl7* | 1 | GGCCCGGGTTCGATTCCCGGCCGATGCAGTCTCAAAGAGGAGCTCGCCGTTTCAGAGCTATGCTGGAAA |
| *mthl7* | 2 | CGAAACTATACAGCATACAATGCACCAGCCGGGAATCGAACC |
| *mthl7* | 3 | TTGTATGCTGTATAGTTTCGGTTTCAGAGCTATGCTGGAAAC |
| *mthl7* | 4 | TTTAACTTGCTATTTCTAGCTCTAAAACATAGTTAGGGTCGACTCGATTGCACCAGCCGGGAATCGAAC |
| *mthl12* | 1 | GGCCCGGGTTCGATTCCCGGCCGATGCAGAAATTCCTGCCAACCTGACGTTTCAGAGCTATGCTGGAAA |
| *mthl12* | 2 | ACATGGTTAGTTTAATTTCATGCACCAGCCGGGAATCGAACC |
| *mthl12* | 3 | TGAAATTAAACTAACCATGTGTTTCAGAGCTATGCTGGAAAC |
| *mthl12* | 4 | TTTAACTTGCTATTTCTAGCTCTAAAACAGTAGATTTCGGGATAAAGATGCACCAGCCGGGAATCGAAC |
| *mth* | 1 | GGCCCGGGTTCGATTCCCGGCCGATGCATTTCGACACTGTCGATATTTGTTTCAGAGCTATGCTGGAAA |
| *mth* | 2 | TGATATGGTCGTGAGGGCAGTGCACCAGCCGGGAATCGAACC |
| *mth* | 3 | CTGCCCTCACGACCATATCAGTTTCAGAGCTATGCTGGAAAC |
| *mth* | 4 | TTTAACTTGCTATTTCTAGCTCTAAAACCTTGCGAAGAGTCACACGGTTGCACCAGCCGGGAATCGAAC |
| *mthl2* | 1 | GGCCCGGGTTCGATTCCCGGCCGATGCAGATACCGTTGATATCTCGGAGTTTCAGAGCTATGCTGGAAA |
| *mthl2* | 2 | TCGTCATGTTGGCGTAGCATTGCACCAGCCGGGAATCGAACC |
| *mthl2* | 3 | ATGCTACGCCAACATGACGAGTTTCAGAGCTATGCTGGAAAC |
| *mthl2* | 4 | TTTAACTTGCTATTTCTAGCTCTAAAACATAGTGCCGACTGATGATAGTGCACCAGCCGGGAATCGAAC |
| *mthl3* | 1 | GGCCCGGGTTCGATTCCCGGCCGATGCATCTTACAATTTATACGCCTGGTTTCAGAGCTATGCTGGAAA |
| *mthl3* | 2 | GAGGGTAAGACTGATGACAGTGCACCAGCCGGGAATCGAACC |
| *mthl3* | 3 | CTGTCATCAGTCTTACCCTCGTTTCAGAGCTATGCTGGAAAC |
| *mthl3* | 4 | TTTAACTTGCTATTTCTAGCTCTAAAACAATTGAGCCTATTGGTTGTTTGCACCAGCCGGGAATCGAAC |
| *mthl4* | 1 | GGCCCGGGTTCGATTCCCGGCCGATGCACTACTTGGCCTCTTTGTTCTGTTTCAGAGCTATGCTGGAAA |
| *mthl4* | 2 | CAGGCGTATAAGTTGTATGCTGCACCAGCCGGGAATCGAACC |
| *mthl4* | 3 | GCATACAACTTATACGCCTGGTTTCAGAGCTATGCTGGAAAC |
| *mthl4* | 4 | TTTAACTTGCTATTTCTAGCTCTAAAACAAAGGACAAACATTATTATGTGCACCAGCCGGGAATCGAAC |
| *ninaE* | 1 | GGCCCGGGTTCGATTCCCGGCCGATGCACTGGAACCAGTTCCCCGCCAGTTTCAGAGCTATGCTGGAAA |
| *ninaE* | 2 | AGCCCAGTCCGGCGTATATGTGCACCAGCCGGGAATCGAACC |
| *ninaE* | 3 | CATATACGCCGGACTGGGCTGTTTCAGAGCTATGCTGGAAAC |
| *ninaE* | 4 | TTTAACTTGCTATTTCTAGCTCTAAAACCACGAGGTCAGGTTACCCTCTGCACCAGCCGGGAATCGAAC |
| *rh2* | 1 | GGCCCGGGTTCGATTCCCGGCCGATGCAGCCGGAGACGCCATTCGACCGTTTCAGAGCTATGCTGGAAA |
| *rh2* | 2 | AGTCTCGTAGTAGAAGTTAATGCACCAGCCGGGAATCGAACC |

| | | |
|---|---|---|
| *rh2* | 3 | TTAACTTCTACTACGAGACTGTTTCAGAGCTATGCTGGAAAC |
| *rh2* | 4 | TTTAACTTGCTATTTCTAGCTCTAAAACCAGGGATTTCACATTCATCTTGCACCAGCCGGGAATCGAAC |
| *rh3* | 1 | GGCCCGGGTTCGATTCCCGGCCGATGCAGCCGGAGACGCCATTCGACCGTTTCAGAGCTATGCTGGAAA |
| *rh3* | 2 | TGGTGGAAGCTATTGTAGATTGCACCAGCCGGGAATCGAACC |
| *rh3* | 3 | ATCTACAATAGCTTCCACCAGTTTCAGAGCTATGCTGGAAAC |
| *rh3* | 4 | TTTAACTTGCTATTTCTAGCTCTAAAACCAAAGAGTCGCGTATCGAAGTGCACCAGCCGGGAATCGAAC |
| *rh4* | 1 | GGCCCGGGTTCGATTCCCGGCCGATGCATTCAATACATTCCGGAGCACGTTTCAGAGCTATGCTGGAAA |
| *rh4* | 2 | AGGGCGAATCCTCGATGGAATGCACCAGCCGGGAATCGAACC |
| *rh4* | 3 | TTCCATCGAGGATTCGCCCTGTTTCAGAGCTATGCTGGAAAC |
| *rh4* | 4 | TTTAACTTGCTATTTCTAGCTCTAAAACCGATCTGCGAGTAGTAGTAATGCACCAGCCGGGAATCGAAC |
| *rh5* | 1 | GGCCCGGGTTCGATTCCCGGCCGATGCATGCACATCAACGGCCCAAGTGTTTCAGAGCTATGCTGGAAA |
| *rh5* | 2 | CCGTTCAAAGCGTATATATCTGCACCAGCCGGGAATCGAACC |
| *rh5* | 3 | GATATATACGCTTTGAACGGGTTTCAGAGCTATGCTGGAAAC |
| *rh5* | 4 | TTTAACTTGCTATTTCTAGCTCTAAAACTGTTGGTCAGGTAATCGAAGTGCACCAGCCGGGAATCGAAC |
| *rh6* | 1 | GGCCCGGGTTCGATTCCCGGCCGATGCACCATTGGAGCCCATGTGGTTGTTTCAGAGCTATGCTGGAAA |
| *rh6* | 2 | TATAATTCGCACAGGAAGGGTGCACCAGCCGGGAATCGAACC |
| *rh6* | 3 | CCCTTCCTGTGCGAATTATAGTTTCAGAGCTATGCTGGAAAC |
| *rh6* | 4 | TTTAACTTGCTATTTCTAGCTCTAAAACCGTAGTTGATGATGGTATACTGCACCAGCCGGGAATCGAAC |
| *rh7* | 1 | GGCCCGGGTTCGATTCCCGGCCGATGCAGAATCATCCGCTGTGAATGTGTTTCAGAGCTATGCTGGAAA |
| *rh7* | 2 | TGTAGTGAATGTAGCTGGGGTGCACCAGCCGGGAATCGAACC |
| *rh7* | 3 | CCCCAGCTACATTCACTACAGTTTCAGAGCTATGCTGGAAAC |
| *rh7* | 4 | TTTAACTTGCTATTTCTAGCTCTAAAACATCCAAAGCGATGGCGGTGATGCACCAGCCGGGAATCGAAC |
| *mGluR* | 1 | GGCCCGGGTTCGATTCCCGGCCGATGCATCAGTGTCTGTTTCTCTGCCGTTTCAGAGCTATGCTGGAAA |
| *mGluR* | 2 | CGAGCGTTAGGCTTCTTTTGTGCACCAGCCGGGAATCGAACC |
| *mGluR* | 3 | CAAAAGAAGCCTAACGCTCGGTTTCAGAGCTATGCTGGAAAC |
| *mGluR* | 4 | TTTAACTTGCTATTTCTAGCTCTAAAACTGTCTCAGTCGTTTGGTCGCTGCACCAGCCGGGAATCGAAC |
| *mtt* | 1 | GGCCCGGGTTCGATTCCCGGCCGATGCACATCAAGCTCACGTTGGTAAGTTTCAGAGCTATGCTGGAAA |
| *mtt* | 2 | CCCCCGAGATGACCTTGCGCTGCACCAGCCGGGAATCGAACC |
| *mtt* | 3 | GCGCAAGGTCATCTCGGGGGGTTTCAGAGCTATGCTGGAAAC |
| *mtt* | 4 | TTTAACTTGCTATTTCTAGCTCTAAAACCGGCTTCGTCAGCAGCTTCTTGCACCAGCCGGGAATCGAAC |
| *GABA-B-R1* | 1 | GGCCCGGGTTCGATTCCCGGCCGATGCAATCGCCTCGCCGCACCTGCAGTTTCAGAGCTATGCTGGAAA |
| *GABA-B-R1* | 2 | GGCGTCGCAAATTGCGCACGTGCACCAGCCGGGAATCGAACC |

| | | |
|---|---|---|
| *GABA-B-R1* | 3 | CGTGCGCAATTTGCGACGCCGTTTCAGAGCTATGCTGGAAAC |
| *GABA-B-R1* | 4 | TTTAACTTGCTATTTCTAGCTCTAAAACGTGGATAGTAACCAAGCCCGTGCACCAGCCGGGAATCGAAC |
| *GABA-B-R2* | 1 | GGCCCGGGTTCGATTCCCGGCCGATGCAGAAAACTCCTACACCGGTCGGTTTCAGAGCTATGCTGGAAA |
| *GABA-B-R2* | 2 | GGGCAGCTGCCCAGATACCATGCACCAGCCGGGAATCGAACC |
| *GABA-B-R2* | 3 | TGGTATCTGGGCAGCTGCCCGTTTCAGAGCTATGCTGGAAAC |
| *GABA-B-R2* | 4 | TTTAACTTGCTATTTCTAGCTCTAAAACTTCGATCCGATAGCACCAGGTGCACCAGCCGGGAATCGAAC |
| *GABA-B-R3* | 1 | GGCCCGGGTTCGATTCCCGGCCGATGCACCGGTCCAAGGGACCAGATAGTTTCAGAGCTATGCTGGAAA |
| *GABA-B-R3* | 2 | CGTCGAGGGCTGTGTGTAGATGCACCAGCCGGGAATCGAACC |
| *GABA-B-R3* | 3 | TCTACACACAGCCCTCGACGGTTTCAGAGCTATGCTGGAAAC |
| *GABA-B-R3* | 4 | TTTAACTTGCTATTTCTAGCTCTAAAACGTCGCTCAACGCCGGCGATGTGCACCAGCCGGGAATCGAAC |
| *fz* | 1 | GGCCCGGGTTCGATTCCCGGCCGATGCATTTACCCACCCTGATACAGGGTTTCAGAGCTATGCTGGAAA |
| *fz* | 2 | TTCTCTCTCCGGGAAGAACATGCACCAGCCGGGAATCGAACC |
| *fz* | 3 | TGTTCTTCCCGGAGAGAGAAGTTTCAGAGCTATGCTGGAAAC |
| *fz* | 4 | TTTAACTTGCTATTTCTAGCTCTAAAACCGTAGGCACATCCAACTACCTGCACCAGCCGGGAATCGAAC |
| *fz2* | 1 | GGCCCGGGTTCGATTCCCGGCCGATGCACGGAGTGCCAGTCATACCCAGTTTCAGAGCTATGCTGGAAA |
| *fz2* | 2 | TCCCCAGGAAGATGAGTGGGTGCACCAGCCGGGAATCGAACC |
| *fz2* | 3 | CCCACTCATCTTCCTGGGGAGTTTCAGAGCTATGCTGGAAAC |
| *fz2* | 4 | TTTAACTTGCTATTTCTAGCTCTAAAACTGTAGAGCACCGAGAAGATGTGCACCAGCCGGGAATCGAAC |
| *fz3* | 1 | GGCCCGGGTTCGATTCCCGGCCGATGCACTGGTGCCACTAATAGAATCGTTTCAGAGCTATGCTGGAAA |
| *fz3* | 2 | AGTCAGAGGTGGGAGATGCTTGCACCAGCCGGGAATCGAACC |
| *fz3* | 3 | AGCATCTCCCACCTCTGACTGTTTCAGAGCTATGCTGGAAAC |
| *fz3* | 4 | TTTAACTTGCTATTTCTAGCTCTAAAACGAACGTGGAACAGCCCGGACTGCACCAGCCGGGAATCGAAC |
| *fz4* | 1 | GGCCCGGGTTCGATTCCCGGCCGATGCACGCCAGAAATCCCCGCCTTCGTTTCAGAGCTATGCTGGAAA |
| *fz4* | 2 | CATGGCCCTTGGAGTCCGGATGCACCAGCCGGGAATCGAACC |
| *fz4* | 3 | TCCGGACTCCAAGGGCCATGGTTTCAGAGCTATGCTGGAAAC |
| *fz4* | 4 | TTTAACTTGCTATTTCTAGCTCTAAAACAGCATCAGTTCCATGAAGGCTGCACCAGCCGGGAATCGAAC |
| *smo* | 1 | GGCCCGGGTTCGATTCCCGGCCGATGCATGAAGCACGTGCCCAAATGTGTTTCAGAGCTATGCTGGAAA |
| *smo* | 2 | CGCCGTTGATTTTTTCACACTGCACCAGCCGGGAATCGAACC |
| *smo* | 3 | GTGTGAAAAAATCAACGGCGGTTTCAGAGCTATGCTGGAAAC |
| *smo* | 4 | TTTAACTTGCTATTTCTAGCTCTAAAACGCACAAAGATCACTATGCAATGCACCAGCCGGGAATCGAAC |
| *CG15744* | 1 | GGCCCGGGTTCGATTCCCGGCCGATGCAGCGACGTCGACCGCAGCAGAGTTTCAGAGCTATGCTGGAAA |
| *CG15744* | 2 | AAGGTGTCTTTGTCCAGTTCTGCACCAGCCGGGAATCGAACC |

| | | |
|---|---|---|
| *CG15744* | 3 | GAACTGGACAAAGACACCTTGTTTCAGAGCTATGCTGGAAAC |
| *CG15744* | 4 | TTTAACTTGCTATTTCTAGCTCTAAAACAATTTAGCCACTCACCCGATTGCACCAGCCGGGAATCGAAC |
| *smog* | 1 | GGCCCGGGTTCGATTCCCGGCCGATGCAAACATTATCCGAATCCACTGTTTCAGAGCTATGCTGGAAA |
| *smog* | 2 | CACGTGTCGGCTCATCGATGTGCACCAGCCGGGAATCGAACC |
| *smog* | 3 | CATCGATGAGCCGACACGTGGTTTCAGAGCTATGCTGGAAAC |
| *smog* | 4 | TTTAACTTGCTATTTCTAGCTCTAAAACTCCGTGCTGTTCCAGGCCGCTGCACCAGCCGGGAATCGAAC |
| *CG31760* | 1 | GGCCCGGGTTCGATTCCCGGCCGATGCACGGCGGCGAAAGGATCAAAGGTTTCAGAGCTATGCTGGAAA |
| *CG31760* | 2 | CAAGTTCTCGGTGGCAATGTTGCACCAGCCGGGAATCGAACC |
| *CG31760* | 3 | ACATTGCCACCGAGAACTTGGTTTCAGAGCTATGCTGGAAAC |
| *CG31760* | 4 | TTTAACTTGCTATTTCTAGCTCTAAAACCTCCATTCGTTAGATTCAGGTGCACCAGCCGGGAATCGAAC |
| *CG12290* | 1 | GGCCCGGGTTCGATTCCCGGCCGATGCATTTGGATAACGCCTGGCTTGGTTTCAGAGCTATGCTGGAAA |
| *CG12290* | 2 | TCTCCGCGCTCATCGATGGTTGCACCAGCCGGGAATCGAACC |
| *CG12290* | 3 | ACCATCGATGAGCGCGGAGAGTTTCAGAGCTATGCTGGAAAC |
| *CG12290* | 4 | TTTAACTTGCTATTTCTAGCTCTAAAACCGGCTGTTGGCGCTGATCAATGCACCAGCCGGGAATCGAAC |
| *CG32447* | 1 | GGCCCGGGTTCGATTCCCGGCCGATGCAATCGGATCGCAACGCCTACCGTTTCAGAGCTATGCTGGAAA |
| *CG32447* | 2 | TCATGAGCATGGTGAGGGTGTGCACCAGCCGGGAATCGAACC |
| *CG32447* | 3 | CACCCTCACCATGCTCATGAGTTTCAGAGCTATGCTGGAAAC |
| *CG32447* | 4 | TTTAACTTGCTATTTCTAGCTCTAAAACCCGCTATCCGGGTGTAGATCTGCACCAGCCGGGAATCGAAC |
| *cg44153* | 1 | GGCCCGGGTTCGATTCCCGGCCGATGCACCGCCGCCACCCTTTCAGCCGTTTCAGAGCTATGCTGGAAA |
| *cg44153* | 2 | GGGCACGCAGGTGTGTACCATGCACCAGCCGGGAATCGAACC |
| *cg44153* | 3 | TGGTACACACCTGCGTGCCCGTTTCAGAGCTATGCTGGAAAC |
| *cg44153* | 4 | TTTAACTTGCTATTTCTAGCTCTAAAACGCGTAGTCTCGTTTATTTCGTGCACCAGCCGGGAATCGAAC |
| *boss* | 1 | GGCCCGGGTTCGATTCCCGGCCGATGCAACGGAGAAAGCCATTACCAGGTTTCAGAGCTATGCTGGAAA |
| *boss* | 2 | TTGTTGGGGCCCTTGTGGTATGCACCAGCCGGGAATCGAACC |
| *boss* | 3 | TACCACAAGGGCCCCAACAAGTTTCAGAGCTATGCTGGAAAC |
| *boss* | 4 | TTTAACTTGCTATTTCTAGCTCTAAAACATCGAGTACGTCGTGCACTGTGCACCAGCCGGGAATCGAAC |
| *CG32547* | 1 | GGCCCGGGTTCGATTCCCGGCCGATGCATCGCCTGGGACCGGATGCGCGTTTCAGAGCTATGCTGGAAA |
| *CG32547* | 2 | CAGATCCGCGCGATGCTCGTTGCACCAGCCGGGAATCGAACC |
| *CG32547* | 3 | ACGAGCATCGCGCGGATCTGGTTTCAGAGCTATGCTGGAAAC |
| *CG32547* | 4 | TTTAACTTGCTATTTCTAGCTCTAAAACCTTCTCGCGATGCACGTCCATGCACCAGCCGGGAATCGAAC |
| *ru* | 1 | GGCCCGGGTTCGATTCCCGGCCGATGCACTAAGTGGGGCACCGTCACAGTTTCAGAGCTATGCTGGAAA |
| *ru* | 2 | CTGGTGGATCAGCACCCACCTGCACCAGCCGGGAATCGAACC |

| | | |
|---|---|---|
| *ru* | 3 | GGTGGGTGCTGATCCACCAGGTTTCAGAGCTATGCTGGAAAC |
| *ru* | 4 | TTTAACTTGCTATTTCTAGCTCTAAAACGTTCTCTTGAATAGAGGGCATGCACCAGCCGGGAATCGAAC |
| *rho* | 1 | GGCCCGGGTTCGATTCCCGGCCGATGCAGAATGTAAACGAAACCAAGGGTTTCAGAGCTATGCTGGAAA |
| *rho* | 2 | GCCGATAGACCAGCACCGAATGCACCAGCCGGGAATCGAACC |
| *rho* | 3 | TTCGGTGCTGGTCTATCGGCGTTTCAGAGCTATGCTGGAAAC |
| *rho* | 4 | TTTAACTTGCTATTTCTAGCTCTAAAACGGGGACCCTTGGCGAAGGCGTGCACCAGCCGGGAATCGAAC |
| *stet* | 1 | GGCCCGGGTTCGATTCCCGGCCGATGCAGGCCATGGAGATCCTGCCCGGTTTCAGAGCTATGCTGGAAA |
| *stet* | 2 | AGGGTATGGGTCCCCTTGGCTGCACCAGCCGGGAATCGAACC |
| *stet* | 3 | GCCAAGGGGACCCATACCCTGTTTCAGAGCTATGCTGGAAAC |
| *stet* | 4 | TTTAACTTGCTATTTCTAGCTCTAAAACGATAGTTAGGCCAGCAATGGTGCACCAGCCGGGAATCGAAC |
| *rho-4* | 1 | GGCCCGGGTTCGATTCCCGGCCGATGCAGCGACAGGACACCATCGATAGTTTCAGAGCTATGCTGGAAA |
| *rho-4* | 2 | AGATCAGTTCCTTCAGCTCGTGCACCAGCCGGGAATCGAACC |
| *rho-4* | 3 | CGAGCTGAAGGAACTGATCTGTTTCAGAGCTATGCTGGAAAC |
| *rho-4* | 4 | TTTAACTTGCTATTTCTAGCTCTAAAACGCGACGGCACCGGACAAATGTGCACCAGCCGGGAATCGAAC |
| *rho-5* | 1 | GGCCCGGGTTCGATTCCCGGCCGATGCAGCAGCATGAGCTCCACGCGCGTTTCAGAGCTATGCTGGAAA |
| *rho-5* | 2 | GTCCGCAAACGGACCCGGAGTGCACCAGCCGGGAATCGAACC |
| *rho-5* | 3 | CTCCGGGTCCGTTTGCGGACGTTTCAGAGCTATGCTGGAAAC |
| *rho-5* | 4 | TTTAACTTGCTATTTCTAGCTCTAAAACATCACTATAGTTCGAAAGATTGCACCAGCCGGGAATCGAAC |
| *rho-6* | 1 | GGCCCGGGTTCGATTCCCGGCCGATGCAGAGAATTCCACGGATTCGGAGTTTCAGAGCTATGCTGGAAA |
| *rho-6* | 2 | TTCCGGTTTGAAGATCAGTATGCACCAGCCGGGAATCGAACC |
| *rho-6* | 3 | TACTGATCTTCAAACCGGAAGTTTCAGAGCTATGCTGGAAAC |
| *rho-6* | 4 | TTTAACTTGCTATTTCTAGCTCTAAAACGGGCACCCGCCACAATCGCTTGCACCAGCCGGGAATCGAAC |
| *rho-7* | 1 | GGCCCGGGTTCGATTCCCGGCCGATGCACCGGAGCTGGCTACCCCAGGGTTTCAGAGCTATGCTGGAAA |
| *rho-7* | 2 | GTGCCGCCGGATGTCTTGTTTGCACCAGCCGGGAATCGAACC |
| *rho-7* | 3 | AACAAGACATCCGGCGGCACGTTTCAGAGCTATGCTGGAAAC |
| *rho-7* | 4 | TTTAACTTGCTATTTCTAGCTCTAAAACGAACGAGAATCCCTTGAGGATGCACCAGCCGGGAATCGAAC |
| *Lgr3* | 1 | GGCCCGGGTTCGATTCCCGGCCGATGCATACGGCAGGAGCATCGCCGTGTTTCAGAGCTATGCTGGAAA |
| *Lgr3* | 2 | GATTCTCGTCGCTGCTGTCGTGCACCAGCCGGGAATCGAACC |
| *Lgr3* | 3 | CGACAGCAGCGACGAGAATCGTTTCAGAGCTATGCTGGAAAC |
| *Lgr3* | 4 | TTTAACTTGCTATTTCTAGCTCTAAAACCGGGCAGGTGGTGCAGATGGTGCACCAGCCGGGAATCGAAC |
| *Lgr4* | 1 | GGCCCGGGTTCGATTCCCGGCCGATGCAGGAAATTGCGGACGCCACAGGTTTCAGAGCTATGCTGGAAA |
| *Lgr4* | 2 | GATCGTGCTGCGGCAGTCGTTGCACCAGCCGGGAATCGAACC |

| | | |
|---|---|---|
| *Lgr4* | 3 | ACGACTGCCGCAGCACGATCGTTTCAGAGCTATGCTGGAAAC |
| *Lgr4* | 4 | TTTAACTTGCTATTTCTAGCTCTAAAACGTTAGGAGCAGAGAGCTTAGTGCACCAGCCGGGAATCGAAC |
| *rk* | 1 | GGCCCGGGTTCGATTCCCGGCCGATGCAGGAGAACGTGATGCTGCCATGTTTCAGAGCTATGCTGGAAA |
| *rk* | 2 | TGGAACGACTGCGGTGGCAATGCACCAGCCGGGAATCGAACC |
| *rk* | 3 | TTGCCACCGCAGTCGTTCCAGTTTCAGAGCTATGCTGGAAAC |
| *rk* | 4 | TTTAACTTGCTATTTCTAGCTCTAAAACGGCTGCCCCTTATGGCCCAGTGCACCAGCCGGGAATCGAAC |
| *Lgr1* | 1 | GGCCCGGGTTCGATTCCCGGCCGATGCAAGCACCCGAGTCTGTCCCAGGTTTCAGAGCTATGCTGGAAA |
| *Lgr1* | 2 | ACTAACGAGGTGCTGGAGAGTGCACCAGCCGGGAATCGAACC |
| *Lgr1* | 3 | CTCTCCAGCACCTCGTTAGTGTTTCAGAGCTATGCTGGAAAC |
| *Lgr1* | 4 | TTTAACTTGCTATTTCTAGCTCTAAAACACGCGGTGAGTCCAAAGAAATGCACCAGCCGGGAATCGAAC |
| *CG13575* | 1 | GGCCCGGGTTCGATTCCCGGCCGATGCAAGCTCGAGTTGTACCTAGAAGTTTCAGAGCTATGCTGGAAA |
| *CG13575* | 2 | GCACGGGTCGCATCACGGCCTGCACCAGCCGGGAATCGAACC |
| *CG13575* | 3 | GGCCGTGATGCGACCCGTGCGTTTCAGAGCTATGCTGGAAAC |
| *CG13575* | 4 | TTTAACTTGCTATTTCTAGCTCTAAAACGTCGATGGGCTCGGAGTAGGTGCACCAGCCGGGAATCGAAC |
| *CG12796* | 1 | GGCCCGGGTTCGATTCCCGGCCGATGCATCGAAGCTCCCCGGAAGCGTGTTTCAGAGCTATGCTGGAAA |
| *CG12796* | 2 | CACAGAAGTCGGAGACGGCCTGCACCAGCCGGGAATCGAACC |
| *CG12796* | 3 | GGCCGTCTCCGACTTCTGTGGTTTCAGAGCTATGCTGGAAAC |
| *CG12796* | 4 | TTTAACTTGCTATTTCTAGCTCTAAAACCACAGAAGTACGGTAGCCAATGCACCAGCCGGGAATCGAAC |

Table S3: Primers used for targeted genomic sequencing

| Gene | # | Sequence |
|---|---|---|
| *CG15614* | 1 | TCGTCGGCAGCGTCAGATGTGTATAAGAGACAGCCGGTTGGCTAATTGGGCAGG |
| *CG15614* | 2 | GTCTCGTGGGCTCGGAGATGTGTATAAGAGACAGGCCGATGGCGTCATCTTGGG |
| *CG15614* | 3 | TCGTCGGCAGCGTCAGATGTGTATAAGAGACAGCGTGTCTGAGCCTGGATAGATTCATC |
| *CG15614* | 4 | GTCTCGTGGGCTCGGAGATGTGTATAAGAGACAGCTGTGACGTGACAAGTTCCCGAG |
| *CG15614* | 5 | TCGTCGGCAGCGTCAGATGTGTATAAGAGACAGggaaagggagtaagtggccatatgtt |
| *CG15614* | 6 | GTCTCGTGGGCTCGGAGATGTGTATAAGAGACAGCTATTAGCAGCGGGGTTGTTAGCC |
| *CrzR* | 1 | TCGTCGGCAGCGTCAGATGTGTATAAGAGACAGGGTGGTCAGCAATTGGTCCAC |
| *CrzR* | 2 | GTCTCGTGGGCTCGGAGATGTGTATAAGAGACAGCTATTGGTGTGCGCCAGTGAG |
| *CrzR* | 3 | TCGTCGGCAGCGTCAGATGTGTATAAGAGACAGCATTCGTGGAGGAGTTTTACCAGTGC |
| *CrzR* | 4 | GTCTCGTGGGCTCGGAGATGTGTATAAGAGACAGcaggttagtacgggaatgtaaactcc |
| *CrzR* | 5 | TCGTCGGCAGCGTCAGATGTGTATAAGAGACAGCGCCATCTTCTTCTTCGGCATG |
| *CrzR* | 6 | GTCTCGTGGGCTCGGAGATGTGTATAAGAGACAGcgcaaaaacaagtacgaaatgtggtg |
| *mAchR-A* | 1 | TCGTCGGCAGCGTCAGATGTGTATAAGAGACAGGGCTATACGATCCCTATTCGGGAATG |
| *mAchR-A* | 2 | GTCTCGTGGGCTCGGAGATGTGTATAAGAGACAGAGGATATCATCACCATGACATTGCCC |
| *mAchR-A* | 3 | TCGTCGGCAGCGTCAGATGTGTATAAGAGACAGCGGTGGTGAATCACGCCTCTG |
| *mAchR-A* | 4 | GTCTCGTGGGCTCGGAGATGTGTATAAGAGACAGgtcctacGCTTGACTTGCGC |
| *mAchR-A* | 5 | TCGTCGGCAGCGTCAGATGTGTATAAGAGACAGCGTACACCTGCTTCGGCAG |
| *mAchR-A* | 6 | GTCTCGTGGGCTCGGAGATGTGTATAAGAGACAGctacCTGGAGACGGAGCTTTGTAAAG |
| *PDFR* | 1 | TCGTCGGCAGCGTCAGATGTGTATAAGAGACAGCGATGTTCATTGTTCGCTTTGCAGC |
| *PDFR* | 2 | GTCTCGTGGGCTCGGAGATGTGTATAAGAGACAGGATTCAAATGTTCCGGCCGTCGG |
| *PDFR* | 3 | TCGTCGGCAGCGTCAGATGTGTATAAGAGACAGCCTACCACAAACAGGTACCTAAAGCC |
| *PDFR* | 4 | GTCTCGTGGGCTCGGAGATGTGTATAAGAGACAGCTGCGCGTATCTACGCCATG |
| *PDFR* | 5 | TCGTCGGCAGCGTCAGATGTGTATAAGAGACAGGACACCCTAAAAAGTGATTTCAAAGG |
| *PDFR* | 6 | GTCTCGTGGGCTCGGAGATGTGTATAAGAGACAGGCAGGGAAACTATAAGGGCGAACAG |
| *Tre1* | 1 | TCGTCGGCAGCGTCAGATGTGTATAAGAGACAGGCAGACATGCAGATGGATGAACCG |
| *Tre1* | 2 | GTCTCGTGGGCTCGGAGATGTGTATAAGAGACAGGCCTGTGCGGCAATTCAGCATAG |
| *Tre1* | 3 | TCGTCGGCAGCGTCAGATGTGTATAAGAGACAGGCTCGATTCCAACGACAGAGCTG |
| *Tre1* | 4 | GTCTCGTGGGCTCGGAGATGTGTATAAGAGACAGCTGCCAGGGGATTAGGGTGACGGTTG |

| | | |
|---|---|---|
| *Tre1* | 5 | TCGTCGGCAGCGTCAGATGTGTATAAGAGACAGCAGATCGCAGCGGCCAAGGGCTC |
| *Tre1* | 6 | GTCTCGTGGGCTCGGAGATGTGTATAAGAGACAGCGAGGTATTGCGCTCGTCGTCCAC |
